# Engineering strategy and vector library for the rapid generation of modular light-controlled protein-protein interactions

**DOI:** 10.1101/583369

**Authors:** Alexandra-Madelaine Tichy, Elliot J. Gerrard, Julien M.D. Legrand, Robin M. Hobbs, Harald Janovjak

## Abstract

Optogenetics enables the spatio-temporally precise control of cell and animal behaviour. Many optogenetic tools are driven by light-controlled protein-protein-interactions (PPIs) that are repurposed from natural light-sensitive domains (LSDs). Applying light-controlled PPI to new target proteins is challenging because it is difficult to predict whether one the many available LSDs will yield robust light regulation. As a consequence, fusion protein libraries need to be prepared and tested, but methods and platforms to facilitate this process are currently not available. Here, we developed a genetic engineering strategy and vector library for the rapid generation of light-controlled PPIs. The strategy permits fusing a target protein to LSDs efficiently and in two orientations. The public and expandable library contains 29 vectors with blue, green or red light-responsive LSDs many of which have been previously applied *ex vivo* and *in vivo.* We demonstrate the versatility of the approach and the necessity for sampling LSDs by generating light-activated caspase-9 (casp9) enzymes. Collectively, this work provides a new resource for optical regulation of a broad range of target proteins in cell and developmental biology.

## INTRODUCTION

Optogenetics has revolutionized research in neuroscience, cell biology and developmental biology by allowing the ‘remote control’ of cell and animal behaviour with extraordinary precision (1–5). This precision is achieved by utilizing light as a stimulus that offers unique advantages over pharmacological and genetic manipulation strategies. For instance, light permits unparalleled control in time (e.g., to modulate animal behaviour acutely or to target selected developmental or disease stages; Figure 1A) and in space (e.g., to target selected compartments in a cell or selected cells in a tissue; Figure 1B). Also, light can be readily applied and withdrawn given a sufficiently transparent matrix. Finally, light-activated molecular tools can be paired with genetic targeting to allow an even higher level of precision for specific cell types, tissues or developmental stages (6–10).

**Figure 1.**
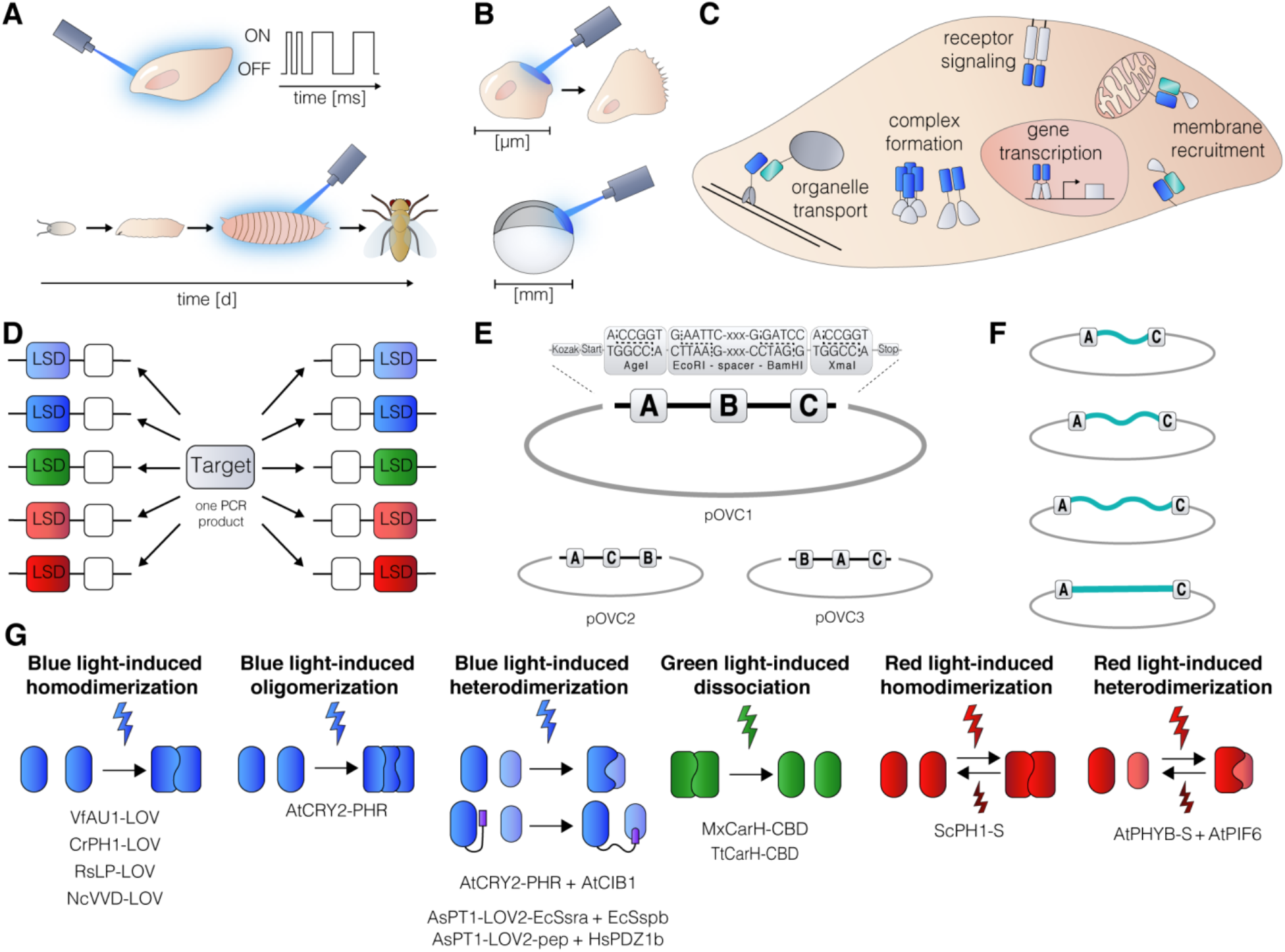
Genetic engineering strategy and vector library for spatio-temporally precise regulation of cell and animal behaviour. (A) The high temporal precision of light can be harnessed to study cellular responses to repetitive and complex inputs and to target specific developmental stages. (B) The high spatial precision of light can be harnessed to selectively activate processes in parts of cells or in specific tissues. (C) Optical control of many cellular processes relies on the regulation of PPIs by incorporating LSDs. (D) Engineering strategy based on a vector library that permits universal target amplicon insertion in a single cloning step (one digest, one ligation). (E) *ABC* and alternative cassettes to realize engineering strategy. Cassettes include Kozak sequence, start codon and stop codon. (F) *ABC* cassettes containing alternative linkers. (G) LSDs and their binding partners included in the library.

Optogenetics first flourished in the hands of neuroscientists that utilized animal and microbial opsins to dissect neural circuits through the bidirectional control of neuronal bioelectrical activity (8,11). More recently and in cell types other than neurons, light control of gene regulation and cellular signalling, together with associated cell behaviours, has emerged (12,13). The optogenetic tools that can regulate cell bioelectricity are fundamentally different from those applied to control biochemical and enzymatic processes. In the former case, ion conducting opsins, such as channelrhodopsin or halorhodopsin, turn neurons on or off by changing their membrane potential through an intrinsic light-gated ion channel or pump activity (7,8,14). In the latter case, a wide range of cellular processes have been rendered light-inducible by using LSDs that do not harbour catalytic activity but regulate intra- or intermolecular binding events (Figure 1C).

LSDs are found in organisms from all domains of life and collectively respond not only to all visible but also to ultraviolet and far-red wavelengths (15–17). Of particular importance in the field of optogenetics are light-oxygen-voltage (LOV) sensing domains and cryptochromes (CRYs) that bind flavins to sense blue light (maximal absorption wavelength (λ_max_) ≈ 450 nm) (18–20) and phytochromes (PHY) that utilize linear tetrapyrroles to sense red (λ_max_ ≈ 660 nm) and far-red (λ_max_ ≈ 720 nm) light (21–23). In addition, green light-sensitive (λ_max_ ≈ 550 nm) cobalamin binding domains (CBDs) that bind vitamin B12 derivatives were applied more recently (24,25). The molecular consequences of photon absorption are either (i) light-induced unmasking of terminal peptides for some LOV domains, (ii) light-induced homodimerization, homooligomerization and heterodimerization with their respective accessory proteins for some LOV domains, CRYs and PHYs, or (iii) light-induced monomerization for some CBDs. These functions have been harnessed in seminal studies to regulate the interactions and activity of diverse target proteins, such as small GTPases, kinases and transcription regulators (6,24,26–34).

The plethora of cellular processes governed by PPIs currently far exceeds the number of available optogenetic tools. This is in part because generating functional fusion proteins of LSDs and target proteins is a non-trivial task. For instance, multiple LSD genes need to be obtained and validated to find a suited domain. Furthermore, the location of the fusion site as well as the length of linkers can be critical parameters that determine fusion protein function (35,36). As a consequence of combinatorial complexity, many genetic constructs need to be generated and tested, and currently no methods or libraries are available to facilitate this process.

Here, we developed a genetic engineering strategy and a vector library for the rapid and modular generation of light-controlled PPIs. The engineering strategy can produce LSD-target protein fusions in several domain orientations and with linkers in a single cloning step (a universal restriction enzyme digest followed by ligation) using inexpensive and readily available materials. The publicly available vector library contains a collection of prominent LSDs that are responsive to blue, green or red light and have been applied in the past *ex vivo* and *in vivo.* The design of the strategy and library allows for easy expansion either with further LSDs, targeting sequences or markers. Using this resource, we generated light-activated casp9 enzymes.

## MATERIALS AND METHODS

### Cassette design

Cassettes were introduced in pcDNA3.1- (Invitrogen/Life Technologies) to generate the vectors named pOVC1-3 (optogenetic vector core 1-3, Figure S4). A XmaI restriction site was removed from the backbone using site-directed mutagenesis (oligonucleotides 1 and 2, Table S2). Inverse polymerase chain reactions (PCR) (oligonucleotides 3 and 4, 5 and 6, and 7 and 8) were applied to remove the vector multiple cloning site and create *ABC* (pOVC1), *ACB* (pOVC2) and *BAC* (pOVC3) cassettes. In the inverse PCR procedure, PCR products were digested with DpnI, digested with EcoRI, XmaI or AgeI (NEB), respectively, ligated for 3 h at room temperature (RT) or overnight at 4°C using T4 ligase (Promega), and propagated in *E.coli* XL 10 Gold cells (Agilent). All cassettes contain Kozak sequences, start codons and stop codons (for backbone ABC, the stop codon was introduced using site-directed mutagenesis in a separate reaction (oligonucleotides 9 and 10)). For linker insertion, backbone pOVC1 was digested using EcoRI and BamHI. Linker fragments were generated by inverse PCR (oligonucleotides 57 and 58) or by annealing and phosphorylating single stranded oligonucleotides (59 to 64). All vector sequences (Table S3) were verified by Sanger sequencing (Micromon, Monash University) and deposited at Addgene.org.

### LSD amplification and vector library

LSDs were amplified using PCR and oligonucleotides with AgeI and/or XmaI restriction site overhangs (oligonucleotides 11 to 34 and 45 to 52). Templates were previously described vectors from our laboratory or obtained from Addgene.org (Table S1). In addition, gene fragments of AtCRY2-PHR, ScPH-1, AsLOV2-EcSsra, EcSSPB micro, AsLOV2-pep and HsPDZ1b were synthesized by a commercial supplier (Integrated DNA Technologies; Table S4). Restriction sites for AgeI and BamHI were removed from ScPH1-S and AtPHYB-S, respectively, as well as XmaI restriction sites from HsFKBP and AtCRY2-PHR using site directed mutagenesis (oligonucleotides 35 to 42). Site-directed mutagenesis was used to create EcSSPB nano (oligonucleotides 65 and 66). PCR products were digested with DpnI and with AgeI, XmaI or AgeI and XmaI depending on oligonucleotide overhangs. Backbone pOVC1 was digested with AgeI or XmaI for insertion into site *A* or *C,* respectively, and phosphatase treated. Backbone and inserts were ligated either for 3 h at RT or overnight at 4°C using T4 ligase (Promega). All vector sequences (Table S5) were verified by Sanger sequencing (LGC Genomics) and deposited at Addgene.org. Note that for future subcloning of the generated genes, universal oligonucleotides can be designed that contain recognition sites for the enzymes AflII, ApaI, AscI, FseI, PacI, PspOMI or SbfI as these are not found in any of the genes.

### Opto-casp9 constructs

The catalytic domain of casp9 (residues 135-416 of UniProt entry P55211) was synthesized (Integrated DNA Technologies; Table S4), amplified by PCR (oligonucleotides 43 and 44) and digested with XmaI. Vectors were digested with XmaI or AgeI, respectively, treated with phosphatase and gel purified. Backbone vectors and casp9 insert were ligated either for 3 h at RT or overnight at 4°C using T4 ligase. Site-directed mutagenesis was used to introduce point substitution C287A into the catalytic domain of casp9 in VfAU1-LOV-casp9, AtCRY2-PHR-casp9 and HsFKBP-casp9 (oligonucleotides 55 and 56). HA-P2A-myc was synthesized as a gene fragment (Integrated DNA Technologies, Table S4), amplified using PCR and restriction site overhangs (oligonucleotides 53 and 54), and inserted into site *B* of VfAU1-LOV-casp9 using EcoRI and BamHI. All vector sequences (Table S6) were verified by Sanger sequencing (Micromon, Monash University) and deposited at Addgene.org.

### Cell culture and transfection

HEK293 cells (Thermo Fisher Scientific; further authenticated by assessing cell morphology and growth rate) were cultured in mycoplasma-free Dulbecco’s modified eagle medium (DMEM, Thermo Fisher Scientific) in a humidified incubator with 5% CO_2_ atmosphere at 37°C. Medium was supplemented with 10% FBS, 100 U/ml penicillin and 0.1 mg/ml streptomycin (Thermo Fisher Scientific). On the day after seeding, cells were transfected in DMEM supplemented with 5% FBS using polyethylenimine (Polysciences). Media was changed after 4 to 6 h and cells were stimulated with light starting 24 h after transfection for the durations specified below and at the intensities specified in the main text.

### Light stimulation of cells

For light stimulation of cells, a tissue culture incubator was equipped with 150 light-emitting diodes (SMD5050-RGB on a LED strip at 3.3 cm spacing). Light intensity was adjusted with a dimmer and measured with a digital power meter (LP1, Sanwa). To obtain light dose curve (Figure S2), one to four layers of neutral density filters (Filter 210, LEE Filters) were used to reduce intensity of selected wells.

### Viability assays

2.5 x 10^4^ HEK293 cells were seeded in each well of white bottom 96-well plates (Costar) and maintained as described above. Cells were transfected with 200 ng vector (casp9, renilla luciferase reporter and empty vector at a ratio of 10:1:9) as described above. Twenty-four h after transfection, cells were either stimulated with blue light (λ ≈ 470 nm), 10 nM of the chemical dimerizer AP20187 (ClonTech Laboratories) or left unstimulated. Light and AP20187 was applied for 7 h. Unstimulated cells were kept in the dark for 6 h before addition of resazurin (Sigma) at a final concentration of 55 μM. After incubation for 1 h, resazurin fluorescence was measured in a plate reader (excitation 540 ±15 nm, emission 590 ± 20 nm, ClarioSTAR, BMGLabtech). Viability was defined as relative fluorescence units compared to a mock transfected control. Immediately after resazurin assays, stimulated and unstimulated cells were washed once with phosphate buffered saline (PBS), lysed and processed with homemade luciferase reporter reagents (24). Luminescence was measured in the plate reader and transcriptional activity was defined as mean raw luminescence values.

### Flow cytometry

5 x 10^5^ HEK293 cells in each well of clear 6-well plates (Costar) were transfected with 2 μg vector (casp9 and empty vector at a ratio of 1:1) as described above. Twenty-four h after transfection, cells were either stimulated with light (blue light, λ ≈ 470 nm, I ≈ 200 μW/cm^2^) or protected from light for 7 h at 37°C. After incubation, cells were collected, washed once with ice-cold PBS (Thermo Fisher Scientific) supplemented with 2% FBS and stained with FITC-AnnexinV/PI Apoptosis Detection Kit according to manufacturer’s instructions (BioLegend). Samples were then run on a LSRFortessa X-20 flow cytometer (BD Biosciences) and data were analysed with FlowJo software (FlowJo).

### Immunoblotting

5 x 10^5^ HEK293 cells in each well of clear 6-well plates (Costar) were transfected with 2 μg vector as described above. Twenty-four h after transfection, cells were washed once with ice-cold PBS and lysed on ice in 180 μl lysis buffer (150 mM NaCl, 1 % TritonX-100, 0.1 % SDS, 0.5 % sodium deoxycholate, 50 mM Tris, complete protease inhibitor (Roche). Lysates were shaken for 30 min at 4°C and centrifuged for 20 min at 12,000xg. 30 μl lysate per lane were separated by SDS-PAGE and electro-blotted onto PVDF membranes. Blots were incubated with primary antibodies (HA-Tag #2367, dilution 1:1000; myc-Tag #2276, dilution 1:1000; Cell Signaling Technology; Vincullin ab129002, dilution 1:10000; Abcam) in blocking solution (5 % BSA in TBS-T) overnight at 4°C. Secondary antibody (goat anti-rabbit IgG(H+L)-HRP conjugate, goat antimouse IgG(H+L)-HRP, dilution 1:10000, Biorad) was applied for 1 h at RT and blots were developed with Carity^™^ Western ECL Substrate (Biorad).

### Statistical analysis

Results were evaluated using Prism (GraphPad). Differences between stimulated and unstimulated samples were analysed using two-tailed T-tests and p values are given in the figure captions. Sample numbers (n) and the number of independent experiments (biological replicates) for each bar are specified in the figure captions, except for Figure 2B-E. In Figure 2B, sample numbers are 14, except for mock (26), HsFKBP (15) and DMSO (13). In Figure 2C, sample numbers are 16, except for mock (25), HsFKBP (19) and DMSO (12). In Figure 2D, sample numbers are 14, except for HsFKBP (26 and 12, dark and light). In Figure 2E, sample numbers are 16, except for mock (28) and HsFKBP (28 and 12, dark and light) and DMSO (12).

**Figure 2.**
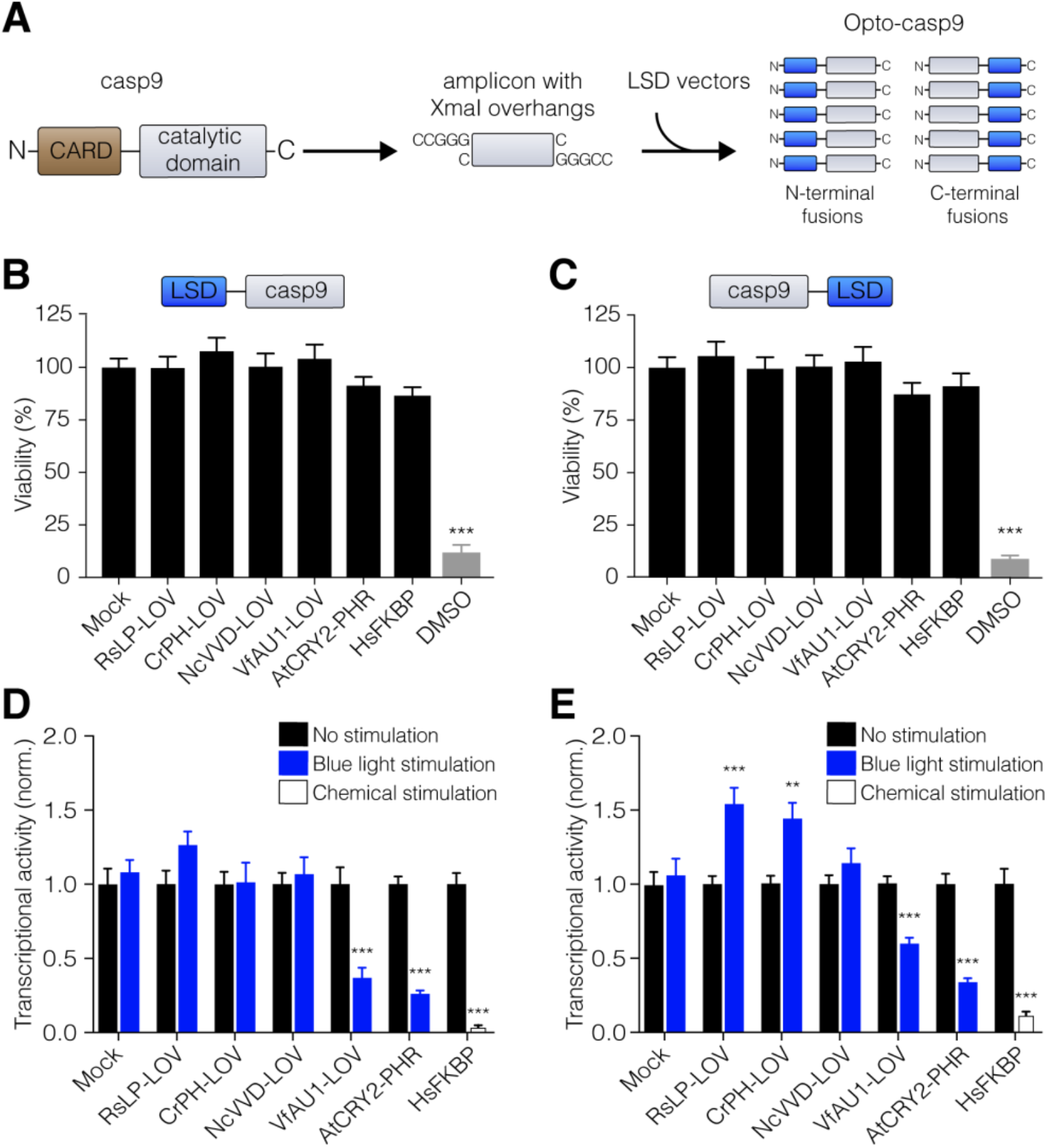
Development of Opto-casp9 enzymes. (A) Procedure to engineer ten Opto-casp9 candidate enzymes starting from one casp9 catalytic domain amplicon and the vector library. (B and C) Viability of cultured human cells transfected with N- (B) and C-terminal (C) fusions of LSDs to casp9 assessed with resazurin. 20% DMSO was employed as a positive control to induce cell death. (D and E) Light-induced cell death for cells transfected with N- (D) and C-terminal (E) fusions of LSDs to casp9 assessed using a luciferase reporter (7 h blue light, λ ≈ 470 nm, I ≈ 200 μW/cm^2^). For B-E: n=12-20 (see Materials and Methods for details), four independent experiments, data shown are mean ± SEM. p<0.001 (**) or p<0.0001 (***). Two-tailed T-test.

## RESULTS AND DISCUSSION

### Efficient genetic engineering strategy

A major challenge in the optical control of PPIs is to achieve functional coupling of LSD oligomerization state changes to activity of target proteins. For most target proteins, it is initially unclear if a suited LSD can be identified and in what orientation LSDs are best attached because steric compatibility and effects on protein folding are difficult to predict. In the majority of previous studies, LSD-target protein fusions were constructed by inserting several LSD genes into vectors that contain the target protein (Figure S1A, top). This approach requires selecting candidate LSDs, obtaining the corresponding genes from collaborators or commercial sources, validating LSD sequences, delineating domain boundaries and preparing amplicons that adapt each LSD to the vector (Figure S1A, bottom). Furthermore, generation of both N- and C-terminal fusion proteins may require additional modification of the vector and/or amplicons. We propose an inverted strategy in which the target protein is inserted into a series of vectors that already contain LSDs (Figure 1D; see below for a comprehensive LSD vector library). The advantages of this strategy are that only a single amplicon of a familiar and available target gene is required and that multiple LSD-target protein fusions can be generated in a simple standardized reaction that is easily parallelized. As a consequence, multiple time-consuming steps that require analysis of sequences and reagents specific to each LSD are not required and the workflow is greatly simplified (Figure S1B).

To achieve this strategy, we designed a modular cloning cassette termed *ABC* that harbours three insertion sites (*A, B* and *C*; Figure 1E). Importantly, sites *A* and *C* contain recognition sequences for restriction enzymes that produce compatible cohesive overhangs (in both cases a CCGG overhang after AgeI or XmaI digestion at site *A* and *C*, respectively; Figure 1E). Consequently, a target protein amplicon flanked by either of these restriction sites in any combination can be inserted into site *A* as well as *C* and thus N- and C-terminally of a LSD (start and stop codons are already contained in the cassette). Site *B* contains recognition sequences for restriction enzymes of different families (EcoRI and BamHI) for incorporation of additional domains (e.g., fluorescent proteins) or epitopes. In order to provide additional flexibility, we engineered *ABC* vectors to include four different flexible or stiff linkers (Figure 1F). We also prepared compatible *ACB* and *BAC* cassettes that permit insertion of flanking targeting sequences or fluorescent proteins in terminal *B* sites. Utilization of single and compatible restriction sites in site *A* and site *C* maximizes the likelihood that target proteins can be inserted without interference from internal restriction sites and minimizes required reagents. Furthermore, restriction enzymes are inexpensive and their application in the cassette retains advanced genetic engineering methods, such as those based on DNA recombination, for transfer of cassettes into other vectors. The promoter region in these vectors can be readily exchanged and promoters and other regulatory sequences are generally part of species-specific or viral vectors. Overall, this genetic engineering strategy permits rapid generation of modular LSD-target protein fusions using readily available reagents.

### LSD vector library

Employing above genetic engineering strategy, we generated 29 vectors that contain one of 11 LSDs or one of five LSD binding partners inserted into site *A* and *C* (Figure 1G). These domains are the photolyase homology region (PHR) domain of plant cryptochrome2 (AtCRY2-PHR of *A. thaliana* (20)), LOV domains of plant, algal and fungal photoreceptors (two modified AsPT1-LOV2 domains of *A. sativa,* CrPH-LOV of *C. reinhardtii,* NcVVD-LOV of *N. crassa,* RsLP-LOV of *R. sphaeroides* and VfAU1-LOV of *V. frigida* (18,19,37–41)), bacterial CBDs (MxCarH-CBD *of M. xanthus* and TtCarH-CBD of *T. thermophilus* (24)), and sensory modules of phytochromes from cyanobacteria and plants (ScPH1-S of *Synechocystis PCC6803* and AtPHYB-S of *A. thaliana* (21,22)). The library also includes binding partners for the heterodimerizing LOV domains, CRY and PHY, which are the minimal proteins EcSspB of *E.coli* with different affinities, HsPDZ1b of *H. sapiens,* AtCIB and AtPIF6 of *A. thaliana* (20,22,39-41); Figure 1G) (sequence information and protein database identifiers can be found in Table S1). Collectively, these vectors provide coverage of methods to induce homodimerization, homooligomerization, heterodimerization with binding partners, or monomerization in response to different wavelengths of light. Many of these domains have been previously utilized *ex vivo* and *in vivo* but the library also contains less frequently applied domains (e.g., CrPH-LOV or RsLP-LOV). Vectors are available with all proteins inserted into the site *A* and separately the site *C* (i.e. N-terminal and C-terminal of the target protein insertion site), except in cases where N-terminal attachment is incompatible with protein function (AsPT1-LOV2 and AtPHYB). In the future, the library is expected to grow as its modular design allows direct expansion with additional LSDs (23,42).

### Light-activated caspase-9

We employed the engineering strategy and vector library to develop a light-induced variant of casp9, an initiator caspase in apoptosis induction. The function of casp9 is mediated by homomeric assembly through the N-terminal caspase recruitment domain (CARD) (43), and casp9 has been rendered inducible by substitution of CARD with orthogonal homodimerization domains (44,45). This work demonstrated that dimerization by an N-terminal domain is sufficient for casp9 activation and resulted in a chemically-induced casp9 (iCasp9) that is employed as a cellular safety and suicide switch (46). To generate casp9 activated by blue light (Opto-casp9), we inserted a casp9 amplicon N-terminally and C-terminally of four LOV domains and AtCRY2-PHR (Figure 2A). We focused on these blue light-sensitive domains because they represent commonly applied optogenetic tools and because their flavin co-factors are ubiquitously available in cells of virtually all organisms. As a control, we employed casp9 fused to an engineered chemical dimerization domain derived from human FK506 binding protein (HsFKBP) analogous to iCasp9. We first tested if these proteins exhibit constitutive activity (i.e. dark activity) by metabolically assessing the viability of human embryonic kidney 293 (HEK293) cells using the fluorescent viability dye resazurin (Figure 2B,C). As constitutive activity was not observed, we next tested if these proteins can be used to induce cell death. To analyse cell death while controlling for transfection efficiency, we co-transfected cells with Opto-casp9 and a genetic viability reporter (*Renilla* luciferase under the control of a constitutive promoter). We chose a luciferase over a fluorescent protein as the reporter gene because of the high signal-to-noise ratio in luminescence detection and to avoid undesired excitation of the reporter by stimulation light. Twenty-four h after transfection, cells were stimulated for 7 h with blue light (λ ≈ 470 nm, intensity (I) = 200 μW/cm^2^) in a tissue culture incubator equipped with light emitting diodes, and luminescence signals were measured subsequently. We found strongly reduced viability for cells that were transfected with casp9 fused to VfAU1-LOV or AtCRY2-PHR domains but not the other domains (Figure 2D,E). To confirm the specificity of the observed effect using VfAU1-LOV-casp9 as an example, we demonstrated that apoptosis increases with increasing light dose (the half maximal effective light dose was 5.5 μW/cm^2^; Figure S2). We further verified that light stimulation resulted in apoptosis using flow cytometry analysis with propidium iodide (PI) and Annexin markers (Figure 3A). For VfAU1-LOV-casp9 and AtCRY2-PHR-casp9 but not for mock transfected cells we observed robust induction of apoptosis (Figure 3B,C). This result demonstrates that by linking a casp9 amplicon to multiple LSDs functional Opto-casp9 enzymes could be quickly designed.

**Figure 3.**
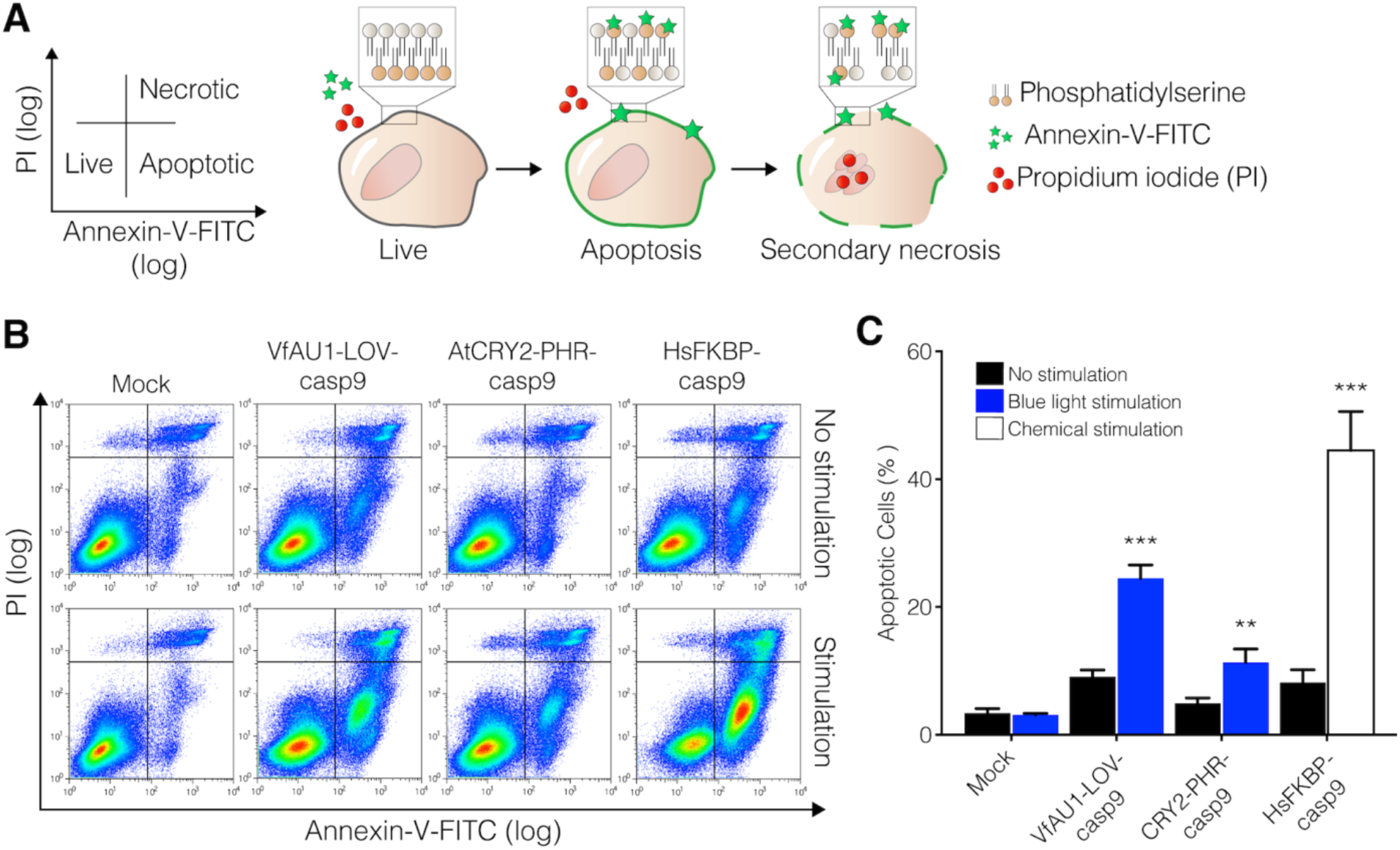
Opto-casp9 induces apoptosis upon light stimulation. (A) Annexin V binding and PI uptake report on apoptotic and necrotic cells. (B) Representative FACS analysis for HEK293 cells that were transfected with VfAU1-LOV-casp9 or AtCRY2-PHR-casp9 and stimulated with light (7 h blue light, λ ≈ 470 nm, I ≈ 200 μW/cm^2^). (C) Quantitative analysis of apoptotic cells (n=3, three independent experiments, data shown are mean ± SEM). p<0.01 (**) or p<0.001 (***). Two-tailed T-test.

### Specificity in light-induced PPIs

The modularity of the genetic engineering strategy provides the possibility to perform additional experiments, such as negative controls and immunodetection, that complement the efficient fusion protein generation demonstrated above. In optogenetics, negative controls typically consist of the application of light to naïve cells or to cells that were transfected with inactivated optogenetic tools (e.g., through loss-of-function mutations). The latter control is required to obtain baseline signals and to ensure that overexpression of LSDs or target proteins does not alter cellular sensitivity. The most commonly applied loss-of-function mutations for inactivation either target photochemically-active LSD residues or residues involved in light-induced conformational changes. However, targeting LSD photochemistry can be incomplete with persistent LSD activation through alternative reaction mechanisms or generation of chemical photoreaction side products (47,48). In addition, because of the diversity in the structures and activation mechanisms of LSDs, generalizable loss-of-function mutations do not exist, and thus negative controls cannot be studied under identical conditions. To address these limitations, we developed a universal inactivation strategy for light-controlled PPIs, which is based on testing the function of constructs in which the LSD and target protein have been uncoupled (e.g., uncoupling of VfAU1-LOV and casp9 should result in a loss in light activation). We realized this strategy by taking advantage of the availability of site *B* in all generated vectors. Into this site, we inserted a self-cleaving peptide sequence of porcine teschovirus-1 2A (P2A) that will effectively dissociate the two domains resulting in a loss of light sensitivity (Figure 4A). As expected for a P2A-modified Opto-casp9, we observed that light-induction of cell death by was abolished completely with self-cleavage effectively producing the same experiment outcome as removal of the catalytic activity of casp9 (Figure 4B, Figure S3). Immunoblotting against epitope tags that flanked the P2A sequence verified cleavage as we only detected the single LSD and casp9 domains but not the full protein (Figure 4C). These results demonstrate a new control strategy that preserves target protein and LSD expression and LSD photochemistry taking advantage of linker and epitope incorporation into site *B*.

**Figure 4.**
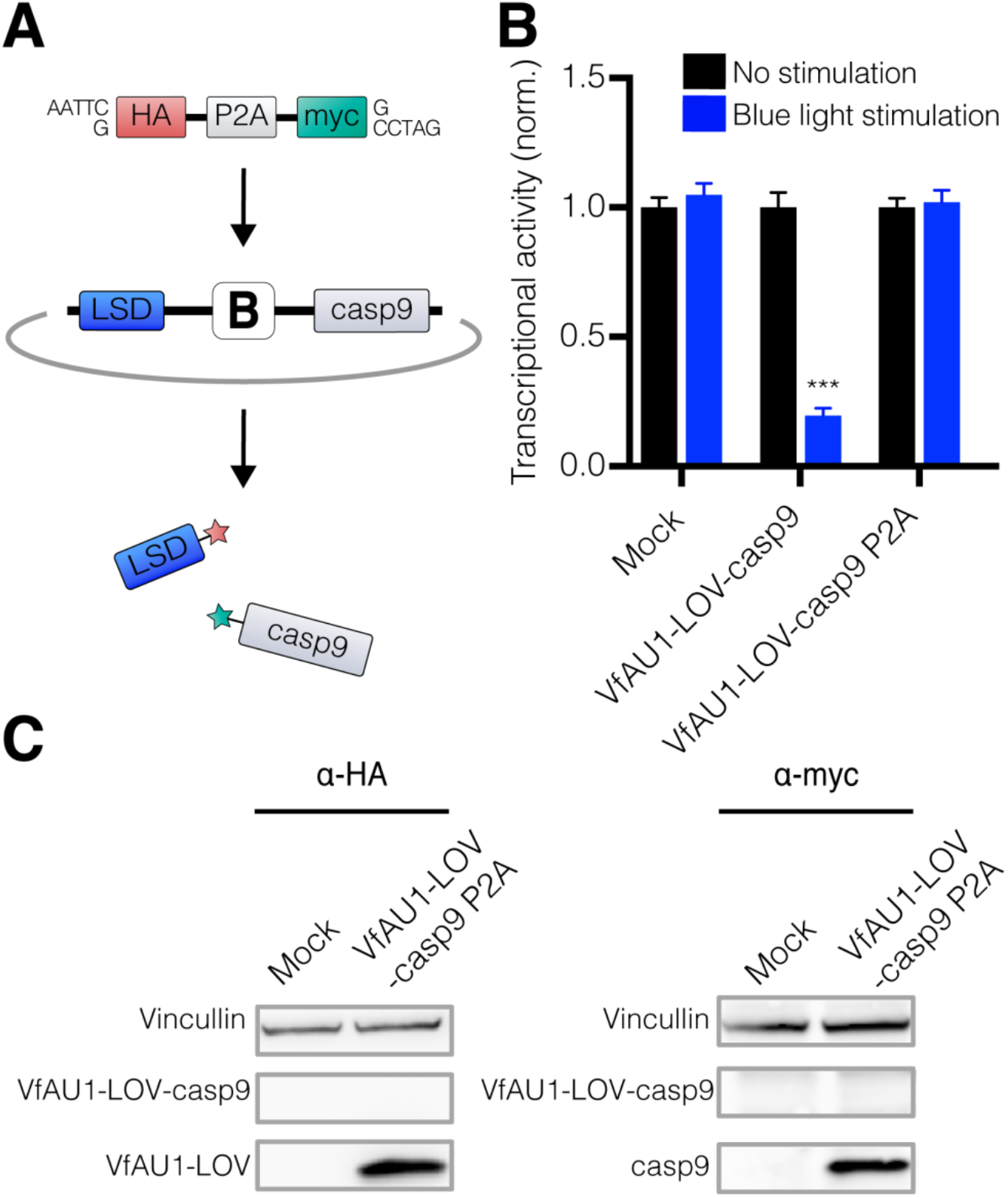
Control experiments with self-cleaving epitope-linker. (A) Incorporation of a P2A sequence results in self-cleavage and separation of LSD and target protein. (B) Light-induced cell death for cells transfected with vectors containing the self-cleaving linker (7 h blue light, λ ≈ 470 nm, I ≈ 200 μW/cm^2^). (C) Immunoblotting to validate efficient cleavage. For B: n=12, four independent experiments, data shown are mean ± SEM. p<0.0001 (***). Two-tailed T-test.

## CONCLUSIONS

Optogenetics is one of few techniques that permits the regulation of cell behaviours with high precision in space and time. We developed a resource for the generation of light-induced PPIs and demonstrated its applicability by engineering Opto-casp9 enzymes. This resource will contribute to the broader use of optogenetics in cell and developmental biology and pave the way to novel optogenetics studies. For instance, experiments on the scale of entire families of LSDs or target proteins require efficient and modular genetic engineering approaches that are now within reach. Opto-casp9 enzymes may provide a test bed for optogenetic hardware development and testing, a process that entails optimization of light parameters (e.g. wavelength, intensity, duration) and culture conditions, because cell death can be assessed with different assays. Finally, the engineering strategy and empty cassettes may also be of use in areas other than optogenetics, such as for the rapid and modular design of fluorescent sensors and protein probes.

## AUTHOR CONTRIBUTIONS

AMT and HJ conceived the study. AMT and HJ designed the engineering strategy and vector library. AMT performed vector library generation and immunoblotting. AMT and EJT designed and constructed Opto-casp9 and performed cell viability assays. JMDL and RH designed flow cytometry analysis. AMT and JMDL performed flow cytometry analysis. All authors interpreted results. AMT and HJ wrote the manuscript. HJ supervised the study.

## ACKNOWLEDGEMENTS

We thank M. De Seram for assistance with experiments, C.P. Heisenberg, E. Gschaider-Reichhart and S. Kainrath for discussions, and C. Tucker and C. Voigt for LSD genes. We acknowledge the facilities, scientific and technical assistance of Micromon and FlowCore at Monash University.

## FUNDING

The Australian Regenerative Medicine Institute is supported by grants from the State Government of Victoria and the Australian Government. The EMBL Australia Partnership Laboratory (EMBL Australia) is supported by the National Collaborative Research Infrastructure Strategy (NCRIS) of the Australian Government. Elliot J. Gerrard was funded through the CSIRO Synthetic Biology Future Science Platform.

## REFERENCES

1. Pastrana, E. (2011) Optogenetics: controlling cell function with light. Nat Meth, 8, 24–25.

2. Reiner, A. and Isacoff, E.Y. (2013) The Brain Prize 2013: The optogenetics revolution. Trends Neurosci, 36, 557–560.

3. Tye, K.M. and Deisseroth, K. (2012) Optogenetic investigation of neural circuits underlying brain disease in animal models. Nat Rev Neurosci, 13, 251–266.

4. Williams, S.C. and Deisseroth, K. (2013) Optogenetics. Proc Natl Acad Sci USA, 110, 16287.

5. Brechun, K.E., Arndt, K.M. and Woolley, G.A. (2017) Strategies for the photo-control of endogenous protein activity. Curr Opin Struct Biol, 45, 53–58.

6. Johnson, H.E., Goyal, Y., Pannucci, N.L., Schupbach, T., Shvartsman, S.Y. and Toettcher, J.E. (2017) The spatiotemporal limits of developmental Erk signaling. Dev Cell, 40, 185–192.

7. Zhang, F., Wang, L.P., Brauner, M., Liewald, J.F., Kay, K., Watzke, N., Wood, P.G., Bamberg, E., Nagel, G., Gottschalk, A. et al. (2007) Multimodal fast optical interrogation of neural circuitry. Nature, 446, 633–639.

8. Boyden, E.S., Zhang, F., Bamberg, E., Nagel, G. and Deisseroth, K. (2005) Millisecond-timescale, genetically targeted optical control of neural activity. Nat Neurosci, 8, 1263–1268.

9. Kowalik, L. and Chen, J.K. (2017) Illuminating developmental biology through photochemistry. Nat Chem Biol, 13, 587–598.

10. Guglielmi, G., Falk, H.J. and De Renzis, S. (2016) Optogenetic control of protein function: From intracellular processes to tissue morphogenesis. Trends Cell Biol, 26, 864–874.

11. Zemelman, B.V., Lee, G.A., Ng, M. and Miesenbock, G. (2002) Selective photostimulation of genetically chARGed neurons. Neuron, 33, 15–22.

12. Toettcher, J.E., Weiner, O.D. and Lim, W.A. (2013) Using optogenetics to interrogate the dynamic control of signal transmission by the Ras/Erk module. Cell, 155, 1422–1434.

13. Muller, K. and Weber, W. (2013) Optogenetic tools for mammalian systems. Mol Biosyst, 9, 596–608.

14. Nagel, G., Szellas, T., Huhn, W., Kateriya, S., Adeishvili, N., Berthold, P., Ollig, D., Hegemann, P. and Bamberg, E. (2003) Channelrhodopsin-2, a directly light-gated cation-selective membrane channel. Proc Natl Acad Sci USA, 100, 13940–13945.

15. Moglich, A., Yang, X., Ayers, R.A. and Moffat, K. (2010) Structure and function of plant photoreceptors. Annu Rev Plant Biol, 61, 21–47.

16. Heijde, M. and Ulm, R. (2012) UV-B photoreceptor-mediated signalling in plants. Trends Plant Sci, 17, 230–237.

17. Ortiz-Guerrero, J.M., Polanco, M.C., Murillo, F.J., Padmanabhan, S. and Elias-Arnanz, M. (2011) Light-dependent gene regulation by a coenzyme B12-based photoreceptor. Proc Natl Acad Sci USA, 108, 7565–7570.

18. Grusch, M., Schelch, K., Riedler, R., Reichhart, E., Differ, C., Berger, W., Ingles-Prieto, A. and Janovjak, H. (2014) Spatio-temporally precise activation of engineered receptor tyrosine kinases by light. EMBO J, 33, 1713–1726.

19. Wang, X., Chen, X. and Yang, Y. (2012) Spatiotemporal control of gene expression by a light-switchable transgene system. Nat Methods, 9, 266–269.

20. Kennedy, M.J., Hughes, R.M., Peteya, L.A., Schwartz, J.W., Ehlers, M.D. and Tucker, C.L. (2010) Rapid blue-light-mediated induction of protein interactions in living cells. Nat Methods, 7, 973–975.

21. Reichhart, E., Ingles-Prieto, A., Tichy, A.M., McKenzie, C. and Janovjak, H. (2016) A phytochrome sensory domain permits receptor activation by red light. Angew Chem Int Ed Engl, 55, 6339–6342.

22. Levskaya, A., Weiner, O.D., Lim, W.A. and Voigt, C.A. (2009) Spatiotemporal control of cell signalling using a light-switchable protein interaction. Nature, 461, 997–1001.

23. Redchuk, T.A., Omelina, E.S., Chernov, K.G. and Verkhusha, V.V. (2017) Near-infrared optogenetic pair for protein regulation and spectral multiplexing. Nat Chem Biol, 13, 633–639.

24. Kainrath, S., Stadler, M., Reichhart, E., Distel, M. and Janovjak, H. (2017) Green-Light-Induced Inactivation of Receptor Signaling Using Cobalamin-Binding Domains. Angew Chem Int Ed Engl, 56, 4608–4611.

25. Jost, M., Fernandez-Zapata, J., Polanco, M.C., Ortiz-Guerrero, J.M., Chen, P.Y., Kang, G., Padmanabhan, S., Elias-Arnanz, M. and Drennan, C.L. (2015) Structural basis for gene regulation by a B12-dependent photoreceptor. Nature, 526, 536–541.

26. Izquierdo, E., Quinkler, T. and De Renzis, S. (2018) Guided morphogenesis through optogenetic activation of Rho signalling during early Drosophila embryogenesis. Nat Commun, 9, 2366.

27. Wang, X., He, L., Wu, Y.I., Hahn, K.M. and Montell, D.J. (2010) Light-mediated activation reveals a key role for Rac in collective guidance of cell movement in vivo. Nat Cell Biol, 12, 591–597.

28. Yoo, S.K., Deng, Q., Cavnar, P.J., Wu, Y.I., Hahn, K.M. and Huttenlocher, A. (2010) Differential regulation of protrusion and polarity by PI3K during neutrophil motility in live zebrafish. Dev Cell, 18, 226–236.

29. Krishnamurthy, V.V., Khamo, J.S., Mei, W., Turgeon, A.J., Ashraf, H.M., Mondal, P., Patel, D. B., Risner, N., Cho, E.E., Yang, J. et al. (2016) Reversible optogenetic control of kinase activity during differentiation and embryonic development. Development, 143, 4085–4094.

30. Sako, K., Pradhan, S.J., Barone, V., Ingles-Prieto, A., Muller, P., Ruprecht, V., Capek, D., Galande, S., Janovjak, H. and Heisenberg, C.P. (2016) Optogenetic control of nodal signaling reveals a temporal pattern of nodal signaling regulating cell fate specification during gastrulation. Cell Rep, 16, 866–877.

31. Reade, A., Motta-Mena, L.B., Gardner, K.H., Stainier, D.Y., Weiner, O.D. and Woo, S. (2017) TAEL: a zebrafish-optimized optogenetic gene expression system with fine spatial and temporal control. Development, 144, 345–355.

32. Beyer, H.M., Juillot, S., Herbst, K., Samodelov, S.L., Muller, K., Schamel, W.W., Romer, W., Schafer, E., Nagy, F., Strahle, U. et al. (2015) Red Light-Regulated Reversible Nuclear Localization of Proteins in Mammalian Cells and Zebrafish. ACS Synth Biol, 4, 951–958.

33. Vopalensky, P., Pralow, S. and Vastenhouw, N.L. (2018) Reduced expression of the Nodal coreceptor Oep causes loss of mesendodermal competence in zebrafish. Development, 145.

34. Barone, V., Lang, M., Krens, S.F.G., Pradhan, S.J., Shamipour, S., Sako, K., Sikora, M., Guet, C.C. and Heisenberg, C.P. (2017) An Effective Feedback Loop between Cell-Cell Contact Duration and Morphogen Signaling Determines Cell Fate. Dev Cell, 43, 198–211 e112.

35. Wu, Y.I., Frey, D., Lungu, O.I., Jaehrig, A., Schlichting, I., Kuhlman, B. and Hahn, K.M. (2009) A genetically encoded photoactivatable Rac controls the motility of living cells. Nature, 461, 104–108.

36. Kawano, F., Okazaki, R., Yazawa, M. and Sato, M. (2016) A photoactivatable Cre-loxP recombination system for optogenetic genome engineering. Nat Chem Biol, 12, 1059–1064.

37. Kutta, R.J., Hofinger, E.S., Preuss, H., Bernhardt, G. and Dick, B. (2008) Blue-light induced interaction of LOV domains from Chlamydomonas reinhardtii. Chembiochem, 9, 1931–1938.

38. Conrad, K.S., Bilwes, A.M. and Crane, B.R. (2013) Light-induced subunit dissociation by a light-oxygen-voltage domain photoreceptor from Rhodobacter sphaeroides. Biochemistry, 52, 378–391.

39. Guntas, G., Hallett, R.A., Zimmerman, S.P., Williams, T., Yumerefendi, H., Bear, J.E. and Kuhlman, B. (2015) Engineering an improved light-induced dimer (iLID) for controlling the localization and activity of signaling proteins. Proc Natl Acad Sci U S A, 112, 112–117.

40. Lungu, O.I., Hallett, R.A., Choi, E.J., Aiken, M.J., Hahn, K.M. and Kuhlman, B. (2012) Designing photoswitchable peptides using the AsLOV2 domain. Chem Biol, 19, 507–517.

41. Strickland, D., Lin, Y., Wagner, E., Hope, C.M., Zayner, J., Antoniou, C., Sosnick, T.R., Weiss, E. L. and Glotzer, M. (2012) TULIPs: tunable, light-controlled interacting protein tags for cell biology. Nat Methods, 9, 379–384.

42. Zhou, X.X., Chung, H.K., Lam, A.J. and Lin, M.Z. (2012) Optical control of protein activity by fluorescent protein domains. Science, 338, 810–814.

43. Renatus, M., Stennicke, H.R., Scott, F.L., Liddington, R.C. and Salvesen, G.S. (2001) Dimer formation drives the activation of the cell death protease caspase 9. Proc Natl Acad Sci U S A, 98, 14250–14255.

44. Nihongaki, Y., Suzuki, H., Kawano, F. and Sato, M. (2014) Genetically engineered photoinducible homodimerization system with improved dimer-forming efficiency. ACS Chem Biol, 9, 617–621.

45. Straathof, K.C., Pule, M.A., Yotnda, P., Dotti, G., Vanin, E.F., Brenner, M.K., Heslop, H.E., Spencer, D.M. and Rooney, C.M. (2005) An inducible caspase 9 safety switch for T-cell therapy. Blood, 105, 4247–4254.

46. Di Stasi, A., Tey, S.K., Dotti, G., Fujita, Y., Kennedy-Nasser, A., Martinez, C., Straathof, K., Liu, E., Durett, A.G., Grilley, B. et al. (2011) Inducible apoptosis as a safety switch for adoptive cell therapy. N Engl J Med, 365, 1673–1683.

47. Yee, E.F., Diensthuber, R.P., Vaidya, A.T., Borbat, P.P., Engelhard, C., Freed, J.H., Bittl, R., Moglich, A. and Crane, B.R. (2015) Signal transduction in light-oxygen-voltage receptors lacking the adduct-forming cysteine residue. Nat Commun, 6, 10079.

48. Shu, X., Lev-Ram, V., Deerinck, T.J., Qi, Y., Ramko, E.B., Davidson, M.W., Jin, Y., Ellisman, M.H. and Tsien, R.Y. (2011) A genetically encoded tag for correlated light and electron microscopy of intact cells, tissues, and organisms. PLoS Biol, 9, e1001041.

