## Supplementary Information for "Engineering strategy and vector library for the rapid generation of modular light-controlled protein-protein interactions"

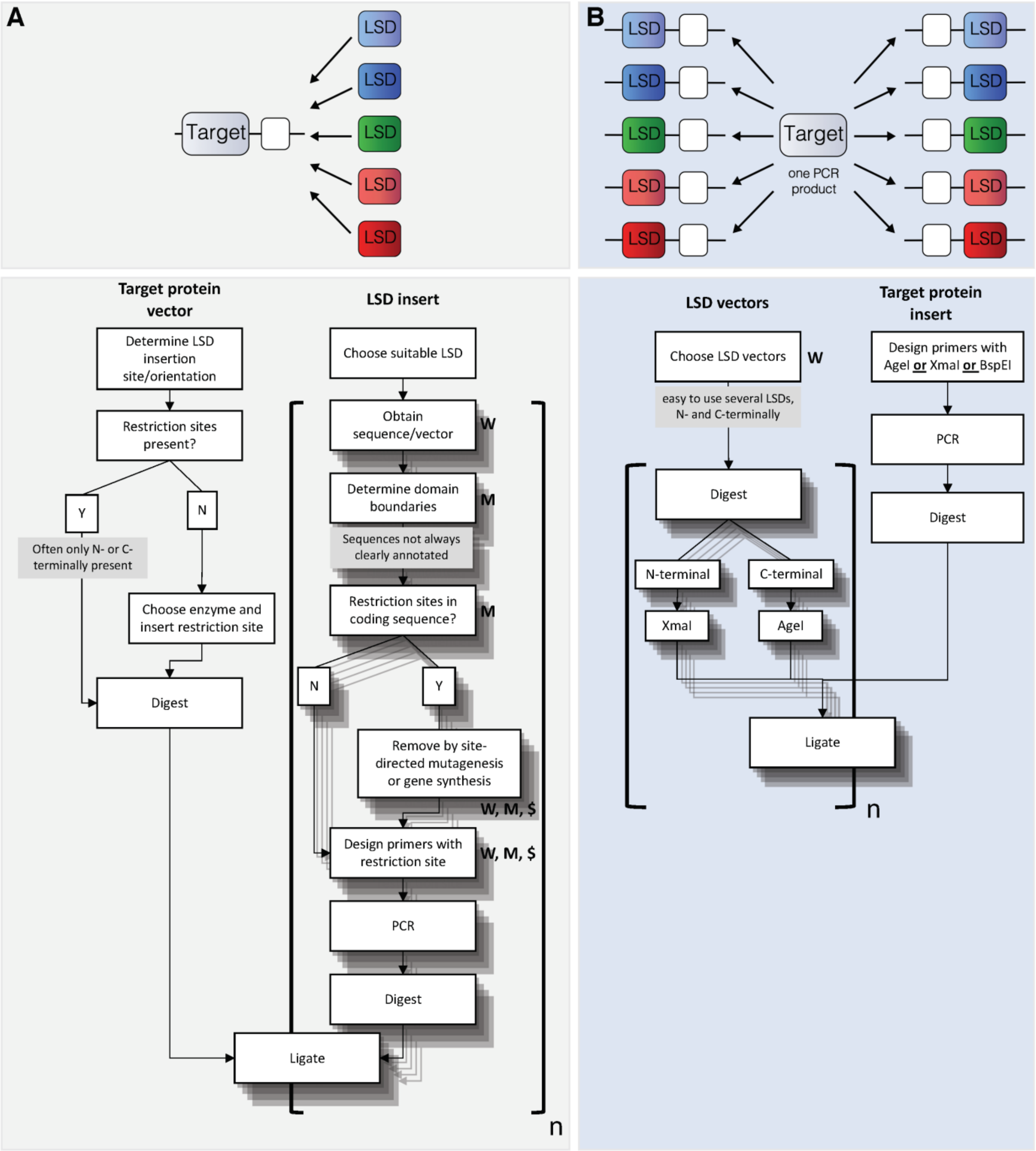


**Supplementary Figure 1.** Optogenetic protein engineering using a conventional workflow in which LSDs are inserted into target vectors (A) or using this genetic engineering strategy and vector library (B). Square brackets highlight steps that need to be repeated for each of the "n" tested LSDs. Within the square brackets, labels denote steps that require manual sequence analysis (M), overnight or longer wait times (W) and reagents ($) that are specific for each LSD.

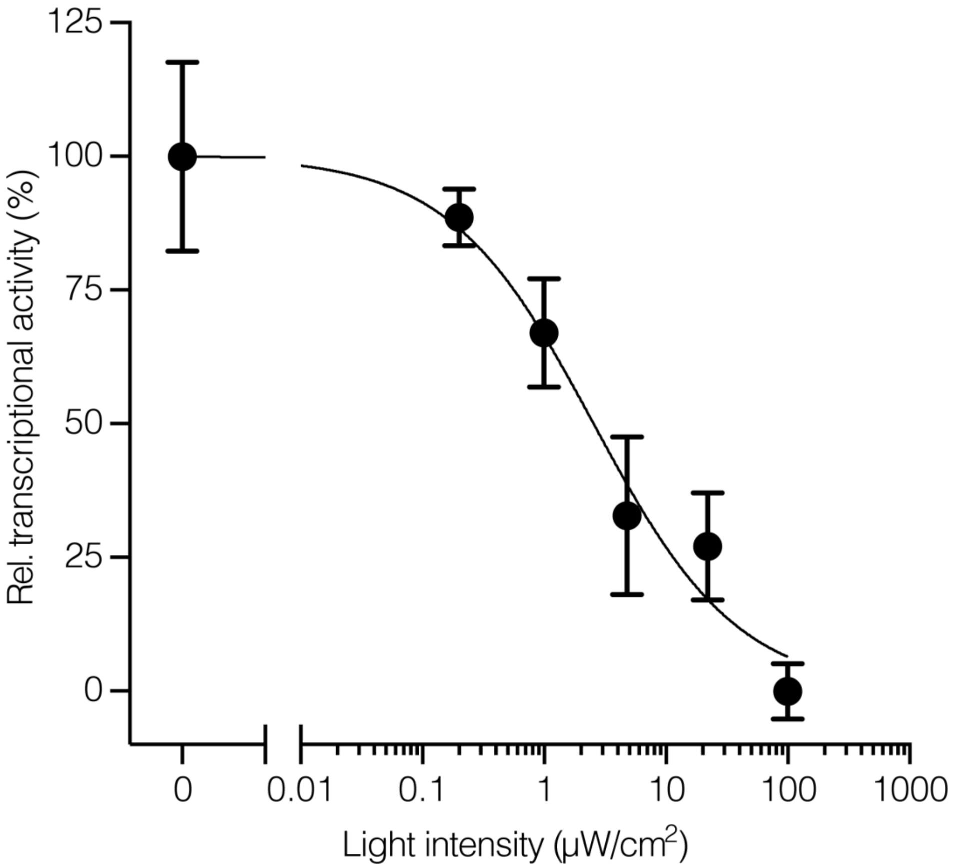


**Supplementary Figure 2.** Viability as a function of light dose (7 h blue light, $\lambda \approx 470$ nm) for cells transfected with VfAU1-LOV-casp9 (one representative experiment in triplicates, data shown are mean $\pm$ SEM, solid line is a sigmoidal dose-response curve fit, EC50 = 5.5 µW/cm$^2$).

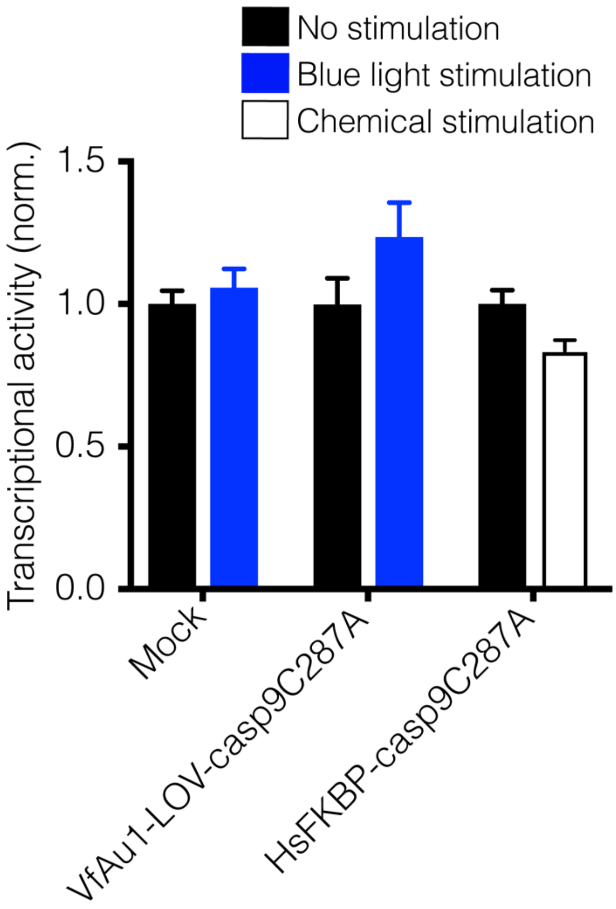


**Supplementary Figure 3.** Viability measurements for cells transfected with Opto-casp9 in which the casp9 active site cysteine was substituted with alanine (C287A) (n=12, four independent experiments, data shown are mean ± SEM; 7 h blue light, $\lambda \approx 470$ nm, $I \approx 200$ μW/cm²).

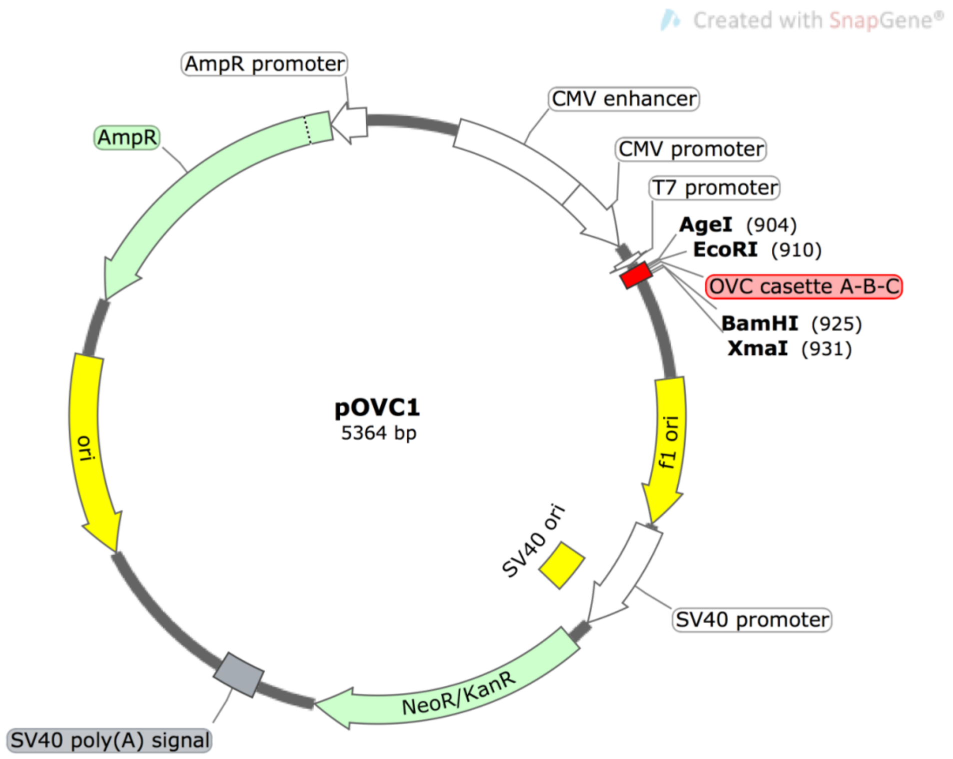


**Supplementary Figure 4. Modified mammalian expression vector.** The original vector pcDNA3.1(-) was modified by removing a XmaI site and inserting one of three cassettes (shown here, pOVC1 with *ABC* cassette: *A*: AgeI, *B*: EcoRI-spacer-BamHI, *C*: XmaI). The cassettes contain a Kozak sequence and start/stop codons. Vector map created using SnapGene.

**Supplementary Table 1.** LSDs and their binding partners included in this study.

| **LSD** | **Cofactor (typ.)** | **Light response** | **Residues (Uniprot identifier)** | **Reference** |
|---|---|---|---|---|
| VfAU1-LOV | Flavin mononucleotide (FMN) | Homodimerization | 204-348 (A8QW55) | (1) |
| CrPH-LOV | FMN | Homodimerization | 16-133 (A8IXU7) | (1) |
| NcVVD-LOV | FMN | Homodimerization | 37-186/Y50W (Q9C3Y6) | (1,2) |
| RsLP-LOV | FMN | Homodimerization | All residues of Protein Data Bank entry 4HJ4 corresponding to genomic position NC_009428 2302851–2303211 | (1,3) |
| AsPT1-LOV2-EcSsra/EcSSPB micro/nano | FMN | Unfolding (heterodimerization, different kinetics with EcSSBP micro or nano) | 404-536 (O49003) | (4) |
| AsPT1-LOV2-pep/HsPDZ1b | FMN | Unfolding (heterodimerization) | 404-537 (O49009) | (5) |
| AtCRY2-PHR | Flavin adenine dinucleotide (FAD) | Oligomerization | 1-498 (Q96524) | (6) |
| AtCRY2-PHR/AtCIB | FAD | Heterodimerization | 1-498 (Q96524)/ 1-170 (Q8GY61) | (7) |
| MxCarH-CBD | Adenosylcobalamin (AdoCbl) | Monomerization | 94-299 (Q50900) | (8) |
| TtCarH-CBD | AdoCbl | Monomerization (irreversible) | 80-284 (Q53W62) | (8) |
| ScPH1-S | Phycocyanobilin (PCB) | Homodimerization | 2-513 (Q55168) | (9) |
| AtPHYB-S/AtPIF | PCB/Phytochromobilin | Heterodimerization | 1-908 (P14713)/ 1-100 (Q8L5W7) | (10) |

**Supplementary Table 2.** Oligonucleotide PCR primers used in this study. Restriction sites are underlined where applicable.

| **Number** | **Sequence** |
|---|---|
| 1 | CTTTTGCAAAAAGCTCTCGGGAGCTTGTATATC |
| 2 | GATATACAAGCTCCCGAGAGCTTTTTGCAAAAG |
| 3 | GATCGAATTCTCAGGCTCTGGATCCCCCGGGCTTAAGTTTAAACCGCTGATCAGCC |
| 4 | GATCGAATTCACCGGTGGCCATGGTCAGCTTGGGTCTCCCTATAGTG |
| 5 | GATCCCCGGGGAATTCTCAGGCTCTGGATCCTAAGTTTAAACCGCTGATCAG |
| 6 | GATCCCCGGGACCGGTGGCCATGGTCAGCTTGGGTCTCCCTATAGTG |
| 7 | GATCGGATCCACCGGTCCCGGGTAAGTTTAAACCGCTGATCAG |
| 8 | GATCGGATCCAGAGCCTGAGAATTCGGCCATGGTCAGCTTGGGTCTCCCTATAGTG |
| 9 | GGATCCCCCGGGTAAGTTTAAACCG |
| 10 | CGGTTTAAACTTACCCGGGGGATCC |
| 11 | AGTCACCGGTCCTGACTACAGTCTCGTGAAGGCT |
| 12 | AGTCCCCGGGCTTTCTGCGCAGCATGTTACTGGT |
| 13 | GATCACCGGTGCAGGACTCAGACATACATTTGTGGTGG |
| 14 | GATCCCCGGGGGCCAGGGCTTTCCCTTCAGTC |
| 15 | GATCCCCGGGCACACTCTCTACGCCCCAGGCG |
| 16 | GATCCCCGGGTTCGGTTTCGCACTGAAAACCCATGCT |
| 17 | GATCACCGGTGCCATGGATCAGAAGCAGTTT |
| 18 | GATCCCCGGGACCCCTTCTTTCCCAGGC |
| 19 | GATCATACCGGTCCACACGCA |
| 20 | GATCATCCCGGGTGTTCCGGCA |
| 21 | GATCATACCGGTCCCGAGGACCT |
| 22 | GATCATCCCGGGGATTGCTTCTTT |
| 23 | GACTCCCGGGGCAACTACTGTTCAACTGTCTGATC |
| 24 | GACTCCCGGGTTCTTCAGCTTGGCGCAGAATCAG |
| 25 | GATCACCGGTAAGATGGACAAGAAAACCATAG |
| 26 | GATCACCGGTTGCAGCACCAATCATGATTTG |
| 27 | GATCACCGGTATGATGTTCTTACCAACCGATTATTG |
| 28 | GATCCCCGGGGTCAACATGTTTATTGCTTTCCAAC |
| 29 | GATCACCGGTGTTTCCGGAGTCGGGGG |
| 30 | GATCCCCGGGGCTCGGGATTTGCAAGAAAC |
| 31 | GATCACCGGTAATGGAGCTATAGGAGGTGACC |
| 32 | GATCCCCGGGACATGAATATAATCCGTTTTCTCC |
| 33 | GATCACCGGTAAACTGGAAGTCGAGGGAGTGC |
| 34 | GATCACCGGTACCGCCAGATTCCAGTTTTAGAAG |
| 35 | GAACTGTACCGGCATCCTGGGCCGTAG |
| 36 | CTACGGCCCAGGATGCCGGTACAGTTC |
| 37 | GAAAGTTGATTCCTCCTGGGACAGAAACAAGC |
| 38 | GCTTGTTTCTGTCCCAGGAGGAATCAACTTTC |
| 39 | GCCTGGTCCCCTGGGTGGAGTAAC |

| | |
|---|---|
| 40 | GTTACTCCACCCAGGGGACCAGGC |
| 41 | GTTAAATCCGGTTTGGATTCATTCCAAGAATACTGG |
| 42 | CCAGTATTCTTGGAATGAATCCAAACCGGATTTAAC |
| 43 | GATCCCGGGGGATTTGGTGATGTCGGTGC |
| 44 | GATCCCCGGGTGATGTTTTAAAGAAAAGTTTTTTCCGG |
| 45 | GATCCCCGGGCTGGCAACCACACTGGAAC |
| 46 | GATCCCCGGGTCCAAAGTAATTTTCGTCGTTCGC |
| 47 | GATCCCCGGGAGCTCCCCGAAACGCCC |
| 48 | GATCCCCGGGACCAATATTCAGCTCGTCATAGATTTC |
| 49 | GATCCCCGGGCCAGAACTTGGATTTAGCATATC |
| 50 | GATCCCCGGGTCTAGAGGTACGGTAGTTAATCGAG |
| 51 | GATCCCCGGGTTGGCTGCTGCACTTGAACG |
| 52 | GATCCCCGGGCACCCAGGTATCCACCGC |
| 53 | GATCGAATTCTACCCATACGATGTTCCAGATTAC |
| 54 | TCGAATTCGAACAAAAACTCATCTCAGAAGAG |
| 55 | CTTTTTCATCCAGGCCGCTGGTGGGGAGCAGAAAG |
| 56 | CTTTCTGCTCCCCACCAGCGGCCTGGATGAAAAAG |
| 57 | GCCGCTTCTTTCGCCGCCGCTTCAGAATTCACCGGTGGCCATG |
| 58 | GGCGAAAGAAGCGGCGGCGAAAGGATCCCCCGGGTAAGTTTA |
| 59 | AATTCGGAAGCTCTAGTGGTG |
| 60 | GATCCACCACTAGAGCTTCCG |
| 61 | AATTCGGAAGCTCTGGCGGATCCAGTGGTG |
| 62 | GATCCACCACTGGATCCGCCAGAGCTTCCG |
| 63 | AATTCGGAAGCAGCTCTTCTGGCGGATCCAGTAGTGGTG |
| 64 | GATCCACCACTACTGGATCCGCCAGAAGAGCTGCTTCCG |
| 65 | CAGTTCAACGCCCGCTTTAAGGGCGTGTC |
| 66 | GACACGCCCTTAAAGCGGGCGTTGAACTG |

**Supplementary Table 3.** Nucleotide sequences of empty and linker-containing cassettes. Green: site *A*, blue: site *B*, red: site *C*. The genes are available through Addgene.org.

| Name | Sequence |
|---|---|
| pOVC1 | ACCATGGCCACCGGTGAATTCTCAGGCTCTGGATCCCCCGGGTAA |
| pOVC2 | ACCATGGCCACCGGTCCCGGGGAATTCTCAGGCTCTGGATCCTAA |
| pOVC3 | ACCATGGCCGAATTCTCAGGCTCTGGATCCACCGGTCCCGGGTAA |
| pOVC1_L1(GSSSG) | ACCATGGCCACCGGTGAATTCGGAAGCTCTAGTGGTGGATCCCCCGGGTAA |
| pOVC1_L2(GSSGGSSG) | ACCATGGCCACCGGTGAATTCGGAAGCTCTGGCGGATCCAGTGGTGGATCCCCCGGGTAA |
| pOVC1_L3(GSSSSGGSSSG) | ACCATGGCCACCGGTGAATTCGGAAGCAGCTCTTCTGGCGGATCCAGTAGTGGTGGATCCCCCGGGTAA |
| pOVC1_L4(EAAAK)$_3$ | ACCATGGCCACCGGTGAATTCGAAGCGGCGGCGAAAGAAGCGGCGGCGAAAGAAGCGGCGGCGAAAGGATCCCCCGGGTAA |

**Supplementary Table 4.** Nucleotide sequences of ordered gene fragments.

| Name | Sequence |
|---|---|
| AtCRY2-PHR | AAGATGGACAAGAAAACCATAGTCTGGTTTCGCCGCGATTTGAGGATAGAGGATAACCCTGCGTTGGCCGCAGCGGCCCACG<br>AAGGCAGCGTGTTCCCCGTGTTCATATGGTGCCCAGAGGAGGAAGGCCAGTTCTACCCAGGTCGCGCTAGTCGCTGGTGGAT<br>GAAACAGTCTCTCGCACATCTCTCCCAATCTTTGAAAGCTCTCGGGTCTGACCTTACGCTGATTAAGACCCACAATACTATT<br>AGTGCAATCTTGGACTGCATCCGCGTTACCGGCGCGACCAAAGTGGTGTTTAATCATCTGTATGACCCGGTAAGTCTGGTAC<br>GCGACCATACAGTTAAGGAGAAGTTGGTTGAAAGGGGAATTTCAGTACAGAGTTATAATGGAGACCTCCTCTACGAACCTTG<br>GGAAATATACTGTGAGAAGGGAAAGCCATTTACATCATTTAACTCATACTGGAAAAAGTGTCTTGACATGAGCATAGAGTCT<br>GTCATGTTGCCCCCGCCGTGGCGGCTTATGCCCATCACCGCGGCCGCCGAAGCTATCTGGGCGTGTAGTATTGAAGAACTTG<br>GCTTGGAGAACGAAGCAGAAAAACCTAGCAATGCACTGTTGACGCGCGCCTGGTCCCCCGGGTGGAGTAACGCAGATAAATT<br>GCTTAACGAGTTCATCGAAAAGCAACTTATCGACTACGCTAAGAATAGCAAAAAAGTGGTGGGCAATTCAACCTCCCTCCTG<br>TCTCCGTACTTGCATTTCGGCGAGATTAGCGTGCGGCACGTCTTCCAATGCGCCCGAATGAAACAAATAATCTGGGCGCGGG<br>ATAAAAACAGTGAGGGTGAAGAAAGCGCAGACTTGTTCCTCCGAGGAATCGGTCTGCGAGAATATAGTCGCTACATTTGTTT<br>CAATTTTCCCTTTACGCATGAGCAGAGCCTCCTTAGTCACTTGCGATTCTTTCCTTGGGACGCAGATGTTGACAAATTTAAA<br>GCATGGCGGCAAGGTAGAACAGGCTACCCATTGGTAGATGCGGGTATGCGAGAACTCTGGGCCACGGGGTGGATGCATAACC<br>GAATCAGGGTAATAGTAAGTAGTTTCGCAGTTAAGTTTCTTTTGCTTCCATGGAAGTGGGGGATGAAGTATTTCTGGGACAC<br>TTTGCTCGATGCGGATCTTGAATGCGATATATTGGGTTGGCAATATATTTCCGGGTCAATCCCTGACGGCCATGAGCTGGAC<br>AGATTGGACAATCCTGCGCTCCAGGGTGCGAAATACGATCCCGAAGGTGAATACATTAGACAATGGCTTCCAGAACTTGCCA<br>GGTTGCCCACGGAATGGATTCACCACCCATGGGACGCCCCTCTTACGGTTTTGAAGGCGAGTGGTGTGGAGCTCGGTACCAA<br>TTACGCAAAACCGATTGTTGACATTGATACCGCGCGGGAGTTGCTGGCTAAAGCCATTTCACGAACCCGGAAAGCCCAAATC<br>ATGATTGGTGCTGCA |
| Casp9 | GATTTGGTGATGTCGGTGCTCTTGAGAGTTTGAGGGGAAATGCAGATTTGGCTTACATCCTGAGCATGGAGCCCTGTGGCCA<br>CTGCCTCATTATCAACAATGTGAACTTCTGCCGTGAGTCCGGGCTCCGCACCCGCACTGGCTCCAACATCGACTGTGAGAAG<br>TTGCGGCGTCGCTTCTCCTCGCTGCATTTCATGGTGGAGGTGAAGGGCGACCTGACTGCCAAGAAAATGGTGCTGGCTTTGC<br>TGGAGCTGGCGCGGCAGGACCACGGTGCTCTGGACTGCTGCGTGGTGGTCATTCTCTCTCACGGCTGTCAGGCCAGCCACCT<br>GCAGTTCCCAGGGGCTGTCTACGGCACAGATGGATGCCCTGTGTCGGTCGAGAAGATTGTGAACATCTTCAATGGGACCAGC<br>TGCCCCAGCCTGGGAGGGAAGCCCAAGCTCTTTTTCATCCAGGCCTGTGGTGGGGAGCAGAAAGACCATGGGTTTGAGGTGG<br>CCTCCACTTCCCCTGAAGACGAGTCCCCTGGCAGTAACCCCGAGCCAGATGCCACCCCGTTCCAGGAAGGTTTGAGGACCTT<br>CGACCAGCTGGACGCCATATCTAGTTTGCCCACACCCAGTGACATCTTTGTGTCCTACTCTACTTTCCCAGGTTTTGTTTCC<br>TGGAGGGACCCCAAGAGTGGCTCCTGGTACGTTGAGACCCTGGACGACATCTTTGAGCAGTGGGCTCACTCTGAAGACCTGC<br>AGTCCCTCCTGCTTAGGGTCGCTAATGCTGTTTCGGTGAAAGGGATTTATAAACAGATGCCTGGTTGCTTTAATTTCCTCCG<br>GAAAAAACTTTTCTTTAAAACATC |
| HA-P2A-MYC | CATGTACCCATACGATGTTCCAGATTACGCTGCGACCAACTTTAGCCTGCTGAAACAGGCGGGCGATGTGGAAGAAAACCCG<br>GGCCCGGAACAAAAACTCATCTCAGAAGAGGATCTGCATG |
| AsPT-LOV2-<br>EcSsra | GATCAGCTGGCAACCACACTGGAACGGATCGAGAAAAATTTCGTGATTACTGATCCGAGACTGCCTGACAACCCAATCATTT<br>TTGCGAGCGATTCCTTCCTGCAGCTGACAGAATATTCTCGGGAAGAGATCCTGGGGCGCAATTGCCGTTTTCTGCAGGGACC<br>CGAGACAGACCGTGCCACTGTTCGGAAAATCAGAGATGCTATTGACAACCAGACTGAAGTGACCGTTCAGCTGATCAATTAT<br>ACCAAGAGCGGCAAGAAGTTCTGGAACGTGTTCCACCTGCAGCCGATGCGCGATTATAAGGGCGACGTCCAGTACTTCATTG<br>GCGTGCAGCTGGATGGCACCGAACGTCTTCATGGCGCCGCTGAGCGTGAGGCGGTCTGCCTGATCAAAAAGACAGCCTTTCA<br>GATTGCTGAGGCAGCGAACGACGAAAATTACTTTGGAGATCAG |
| EcSSPB<br>(micro) | GATCAGAGCTCCCCGAAACGCCCTAAGCTGCTGCGTGAATATTACGATTGGCTGGTTGATAACAGCTTTACCCCATATCTGG<br>TGGTGGATGCCACATACCTGGGCGTGAACGTGCCCGTGGAGTATGTGAAAGACGGTCAGATCGTGCTGAATCTGTCTGCAAG<br>TGCGACCGGCAACCTGCAACTGACAAATGATTTTATCCAGTTCAACGCCCAGTTTAAGGGCGTGTCTCGTGAACTGTATATC<br>CCGATGGGTGCCGCTCTGGCCATTTACGCTCGCGAGAACGGCGATGGTGTGATGTTCGAACCAGAAGAAATCTATGACGAGC<br>TGAATATTGGTGATCAG |

| | |
|---|---|
| AsPT-LOV2-pep | GATCAGTTGGCTGCTGCACTTGAACGTATTGAGAAGAACTTTGTCATTACTGACCCAAGATTGCCAGATAATCCCATTATATTCGCGTCCGATAGTTTCTTGCAGTTGACAGAATATAGCCGTGAAGAAATTTTGGGAAGAAACTGCAGGTTTCTACAAGGTCCTGAAACTGATCGCGCGACAGTGAGAAAAATTAGAGATGCCATAGATAACCAAACAGAGGTCACTGTTCAGCTGATTAATTATACAAAGAGTGGTAAAAAGTTCTGGAACCTCTTTCACTTGCAGCCTATGCGAGATCAGAAGGGAGATGTCCAGTACTTTATTGGGGTTCAGTTGGATGGAACTGAGCATGTCCGAGATGCTGCCGAGAGAGAGGCAGTCATGCTGGCGAAGAAAACTGCAGAAGAGATTGATAAAGCGGTGGATACCTGGGTGGATCAG |
| HsPDZB1 | GATCAGCCAGAACTTGGATTTAGCATATCAGGTGGTGTCGGGGGTAGAGGAAACCCATTCAGACCTGATGATGATGGTATATTTGTAACAAGGGTACAACCTGAAGGACCAGCATCAAAATTACTGCAGCCAGGTGATAAAATTATTCAGGCTAATGGCTACAGTTTTATAAATATTGAACATGGACAAGCAGTGTCCTTGCTAAAAACTTTCCAGAATACAGTTGAACTCATCATTGTACGAGAAGTTGGTAACGGTGCAAAACAAGAGATTCGAGTGAGGGTTGAAAAAGACGGTGGTTCTGGAGGCGTTTCTTCTGTTCCGACCAATCTGGAAGTTGTTGCTGCGACCCCGACTAGCCTGCTGATCAGCTGGGATGCGTATAGGGAGCTTCCGGTTAGTTATTACCGTATCACGTACGGTGAAACAGGTGGTAACTCCCCGGTTCAGGAGTTCACTGTACCTGGTTCCAAGTCTACTGCTACCATCAGCGGCCTGAAACCGGGTGTCGACTATACCATCACTGTATACGCACATTACAACTACCATTACTACTCTAGCCCAATCTCGATTAACTACCGTACCTCTAGAGATCAG |

**Supplementary Table 5.** Nucleotide sequences of cassettes containing LSDs. Green: site *A*, blue: site *B*, red: site *C*. X: Empty site. The genes are available through Addgene.org.

| **Name** | **Sequence** |
|---|---|
| >pOVC1_VfAU1-LOV_X | ACCATGGCCACCGGTCCTGACTACAGTCTCGTGAAGGCTCTGCAAATGGCACAACAGAATTTTGTCATTACAG<br>ACGCCTCCCTCCCAGACAACCCTATCGTCTACGCCAGTAGAGGGTTTCTGACACTGACAGGCTATTCTCTCGA<br>CCAGATCCTGGGCAGGAACTGCAGGTTTCTGCAAGGGCCAGAAACAGACCCAAGAGCTGTGGATAAGATCAGG<br>AATGCCATCACCAAAGGCGTTGATACCAGTGTCTGTCTGCTGAATTATAGACAGGATGGCACAACCTTCTGGA<br>ATCTCTTCTTCGTGGCTGGACTCAGAGATTCTAAGGGCAATATTGTCAACTACGTCGGAGTGCAGTCAAAGGT<br>GAGCGAAGATTATGCCAAGCTGCTGGTCAACGAGCAGAACATTGAGTACAAAGGTGTGCGCACCAGTAACATG<br>CTGCGCAGAAAGCCCGGTGAATTCTCAGGCTCTGGATCCCCCGGGTAA |
| >pOVC1_CrPH-LOV_X | ACCATGGCCACCGGTGCAGGACTCAGACATACATTTGTGGTGGCTGATGCAACACTCCCTGATTGCCCACTGG<br>TCTATGCAAGTGAGGGCTTCTACGCAATGACCGGATATGGACCTGACGAAGTGCTGGGTCACAACTGTAGGTT<br>TCTGCAGGGTGAGGGAACTGACCCCAAGGAAGTGCAGAAAATTCGCGACGCCATCAAGAAGGGTGAGGCTTGT<br>AGTGTGCGCCTCCTGAACTATCGGAAGGACGGCACTCCCTTCTGGAACCTGCTGACAGTCACCCCAATTAAAA<br>CCCCTGATGGCCGCGTGTCCAAGTTTGTCGGCGTGCAGGTGGATGTTACCTCCAAGACTGAAGGGAAAGCCCT<br>GGCCCCCGGTGAATTCTCAGGCTCTGGATCCCCCGGGTAA |
| >pOVC1_NcVVD-LOV_X | ACCATGGCCACCGGGCACACTCTCTACGCCCCAGGCGGGTACGATATTATGGGCTGGCTGATCCAGATCATGA<br>ACAGGCCCAATCCCCAGGTCGAGCTGGGACCCGTGGATACTTCATGTGCACTGATACTGTGCGACCTGAAGCA<br>GAAGGATACACCTATAGTTTACGCTTCAGAAGCCTTTCTGTACATGACAGGGTATTCTAACGCCGAGGTGCTG<br>GGGAGGAACTGTAGGTTCCTCCAGAGTCCCGATGGTATGGTGAAACCTAAGAGTACTCGCAAATATGTGGATA<br>GCAATACTATTAACACCATGAGGAAAGCCATCGACAGAAACGCAGAAGTTCAGGTGGAAGTGGTGAACTTTAA<br>GAAGAACGGCCAGCGGTTCGTGAACTTTCTCACAATGATTCCAGTGCGGGACGAAACCGGGGAGTACCGGTAC<br>AGCATGGGTTTTCAGTGCGAAACCGAACCCGGTGAATTCTCAGGCTCTGGATCCCCCGGGTAA |
| >pOVC1_RsLP-LOV_X | ACCATGGCCACCGGTGCCATGGATCAGAAGCAGTTTGAGAAGATTAGAGCTGTGTTTGACAGGTCAGGGGTCG<br>CACTGACCCTCGTTGACATGTCCCTGCCAGAGCAACCCCTGGTGCTCGCCAACCCTCCATTTCTGAGAATGAC<br>TGGCTATACTGAGGGCCAGATCCTGGGATTCAACTGCAGATTTCTCCAGAGAGGCGACGAAAATGCTCAGGCA<br>CGGGCTGACATCAGAGATGCCCTCAAGCTCGGAAGGGAGCTCCAGGTGGTCCTCCGCAATTACAGAGCCAACG<br>ATGAACCATTTGACAATCTGCTGTTCCTGCACCCTGTCGGTGGCAGACCCGACGCTCCTGACTACTTCCTCGG<br>TTCTCAGTTCGAGCTGGGTAGAAGCGGAAATAGCGAAGAGGCAGCCGCAGCTGGACACGCAGGGGCACTGACT<br>GGGGAGCTCGCCAGAATAGGAACTGTGGCTGCTCGGCTCGAAATGGACAGTCGGAGACATCTGGCACAAGCTG<br>CTGCAGCCCTGGTGAGGGCCTGGGAAAGAAGGGGTCCCGGTGAATTCTCAGGCTCTGGATCCCCCGGGTAA |
| >pOVC1_EcSSPB-nano_X | ACCATGGCCACCGGGAGCTCCCCGAAACGCCCTAAGCTGCTGCGTGAATATTACGATTGGCTGGTTGATAACA<br>GCTTTACCCCATATCTGGTGGTGGATGCCACATACCTGGGCGTGAACGTGCCCGTGGAGTATGTGAAAGACGG<br>TCAGATCGTGCTGAATCTGTCTGCAAGTGCGACCGGCAACCTGCAACTGACAAATGATTTTATCCAGTTCAAC<br>GCCCGCTTTAAGGGCGTGTCTCGTGAACTGTATATCCCGATGGGTGCCGCTCTGGCCATTTACGCTCGCGAGA<br>ACGGCGATGGTGTGATGTTCGAACCAGAAGAAATCTATGACGAGCTGAATATTGGTCCCGGTGAATTCTCAGG<br>CTCTGGATCCCCCGGGTAA |
| >pOVC1_EcSSPB-micro_X | ACCATGGCCACCGGGAGCTCCCCGAAACGCCCTAAGCTGCTGCGTGAATATTACGATTGGCTGGTTGATAACA<br>GCTTTACCCCATATCTGGTGGTGGATGCCACATACCTGGGCGTGAACGTGCCCGTGGAGTATGTGAAAGACGG<br>TCAGATCGTGCTGAATCTGTCTGCAAGTGCGACCGGCAACCTGCAACTGACAAATGATTTTATCCAGTTCAAC<br>GCCCAGTTTAAGGGCGTGTCTCGTGAACTGTATATCCCGATGGGTGCCGCTCTGGCCATTTACGCTCGCGAGA<br>ACGGCGATGGTGTGATGTTCGAACCAGAAGAAATCTATGACGAGCTGAATATTGGTCCCGGTGAATTCTCAGG<br>CTCTGGATCCCCCGGGTAA |
| >pOVC1_HsPDZ1b_X | ACCATGGCCACCGGGCCAGAACTTGGATTTAGCATATCAGGTGGTGTCGGGGGTAGAGGAAACCCATTCAGAC<br>CTGATGATGATGGTATATTTGTAACAAGGGTACAACCTGAAGGACCAGCATCAAAATTACTGCAGCCAGGTGA<br>TAAAATTATTCAGGCTAATGGCTACAGTTTTATAAATATTGAACATGGACAAGCAGTGTCCTTGCTAAAAACT<br>TTCCAGAATACAGTTGAACTCATCATTGTACGAGAAGTTGGTAACGGTGCAAAACAAGAGATTCGAGTGAGGG<br>TTGAAAAAGACGGTGGTTCTGGAGGCGTTTCTTCTGTTCCGACCAATCTGGAAGTTGTTGCTGCGACCCCGAC<br>TAGCCTGCTGATCAGCTGGGATGCGTATAGGGAGCTTCCGGTTAGTTATTACCGTATCACGTACGGTGAAACA<br>GGTGGTAACTCCCCGGTTCAGGAGTTCACTGTACCTGGTTCCAAGTCTACTGCTACCATCAGCGGCCTGAAAC<br>CGGGTGTCGACTATACCATCACTGTATACGCACATTACAACTACCATTACTACTCTAGCCCAATCTCGATTAA<br>CTACCGTACCTCTAGACCCGGTGAATTCTCAGGCTCTGGATCCCCCGGGTAA |
| >pOVC1_MxCarH-CBD_X | ACCATGGCCACCGGTCCACACGCAGAGACTTGGAGGGAATCTATGCTTGCCGCCACACAAGCCTACGACCAGC<br>CTAGAGTATCAGATGTACTGGATGAGGTCCTTGCGGCTCTGCCCCCTCTGAAGGCCTTCGATGAAGTGCTGGC<br>CCCTTTGCTGTGCGATGTCGGAGAGCGGTGGGAGAGCGGAACCCTGACAGTTGCGCAGGAACATCTGGTCTCA<br>CAGATGGTGCGCGCCCGGCTGGTGAGTCTGCTGCACGCGGCACCATTGGGACGCCACAGACATGGCGTTCTCG<br>CCTGTTTCCCAGAGGAGGAGCATGAGATGGGCTTGCTTGGTGCCGCCTTGAGACTCCGCCATCTCGGCGTTAG<br>AGTAACCCTGCTCGGCCAGCGAGTGCCAGCCGAGGACCTCGGGCGAGCAGTGTTGGCCCTGCGCCCGGACTTC<br>GTGGGCCTGTCAACAGTTGCAAGCAGGAGCGCAGAGGACTTCGAGGATACCTTGACCCGACTCCGCCAGGCCC<br>TGCCAAGGGGCCTCCCTGTATGGGTGGGCGGGGCAGCCGCAAGGTCTCATCAGGCCGTGTGCGAGCGCCTGGC |

| | |
|---|---|
| | AGTCCATGTTTTTCAGGGCGAAGAAGATTGGGATAGACTTGCCGGAACACCCGGTGAATTCTCAGGCTCTGGA<br>TCCCCCGGGTAA |
| >pOVC1_TtCarH-CBD_X | ACCATGGCCACCGGTCCCGAGGACCTCGGCACCGGACTCCTCGAAGCACTCTTGAGAGGAGATTTGGCCGGCG<br>CCGAGGCTCTCTTTCGACGGGGGCTTAGGTTTTGGGGACCCGAGGGTATTCTGGAGCACCTGCTCCTGCCTGT<br>GCTTCGGGAGGTGGAGAAGCCTGGCATCGCGGCGAGATCGGTGTGGCCGAGGAGCATCTGGCATCCACATTT<br>CTGCGCGCGAGACTGCAGGAGCTGCTCGACCTCGCCGGGTTCCCACCTGGCCCCCCCGTGCTGGTAACCACGC<br>CACCAGGGGAGAGGCACGAGATCGGCGCAATGTTGGCTGCCTATCACCTGCGAAGGAAGGGCGTGCCAGCGCT<br>GTACTTGGGACCAGACACCCCCCTCCCCGATCTCAGAGCACTTGCGAGACGGCTCGGAGCCGGAGCGGTGGTT<br>CTGTCAGCTTTGCTTTCCGAGCCTCTCAGGGCGTTGCCAGACGGCGCACTGAAAGACTTGGCACCTCGGGTGT<br>TCCTGGGAGGGCAAGGAGCCGGCCCTGAAGAGGCCCGACGGCTCGGGGCCGAGTACATGGAAGATCTGAAGGG<br>ATTGGCCGAAGCACTGTGGCTTCCAAGAGGACCAGAGAAAGAAGCAATCCCCGGTGAATTCTCAGGCTCTGGA<br>TCCCCCGGGTAA |
| >pOVC1_AtCRY2-PHR_X | ACCATGGCCACCGGTAAGATGGACAAGAAAACCATAGTCTGGTTTCGCCGCGATTTGAGGATAGAGGATAACC<br>CTGCGTTGGCCGCAGCGGCCCACGAAGGCAGCGTGTTCCCCGTGTTCATATGGTGCCCAGAGGAGGAAGGCCA<br>GTTCTACCCAGGTCGCGCTAGTCGCTGGTGGATGAAACAGTCTCTCGCACATCTCTCCCAATCTTTGAAAGCT<br>CTCGGGTCTGACCTTACGCTGATTAAGACCCACAATACTATTAGTGCAATCTTGGACTGCATCCGCGTTACCG<br>GCGCGACCAAAGTGGTGTTTAATCATCTGTATGACCCGGTAAGTCTGGTACGCGACCATACAGTTAAGGAGAA<br>GTTGGTTGAAAGGGGAATTTCAGTACAGAGTTATAATGGAGACCTCCTCTACGAACCTTGGGAAATATACTGT<br>GAGAAGGGAAAGCCATTTACATCATTTAACTCATACTGGAAAAAGTGTCTTGACATGAGCATAGAGTCTGTCA<br>TGTTGCCCCCGCCGTGGCGGCTTATGCCCATCACCGCGGCCGCCGAAGCTATCTGGGCGTGTAGTATTGAAGA<br>ACTTGGCTTGGAGAACGAAGCAGAAAAACCTAGCAATGCACTGTTGACGCGCGCCTGGTCCCCTGGGTGGAGT<br>AACGCAGATAAATTGCTTAACGAGTTCATCGAAAAGCAACTTATCGACTACGCTAAGAATAGCAAAAAAGTGG<br>TGGGCAATTCAACCTCCCTCCTGTCTCCGTACTTGCATTTCGGCGAGATTAGCGTGCGGCACGTCTTCCAATG<br>CGCCCGAATGAAACAAATAATCTGGGCGCGGATAAAAACAGTGAGGGTGAAGAAAGCGCAGACTTGTTCCTC<br>CGAGGAATCGGTCTGCGAGAATATAGTCGCTACATTTGTTTCAATTTTCCCTTTACGCATGAGCAGAGCCTCC<br>TTAGTCACTTGCGATTCTTTCCTTGGGACGCAGATGTTGACAAATTTAAAGCATGGCGGCAAGGTAGAACAGG<br>CTACCCATTGGTAGATGCGGGTATGCGAGAACTCTGGGCCACGGGGTGGATGCATAACCGAATCAGGGTAATA<br>GTAAGTAGTTTCGCAGTTAAGTTTCTTTTGCTTCCATGGAAGTGGGGGATGAAGTATTTCTGGGACACTTTGC<br>TCGATGCGGATCTTGAATGCGATATATTGGGTTGGCAATATATTTCCGGGTCAATCCCTGACGGCCATGAGCT<br>GGACAGATTGGACAATCCTGCGCTCCAGGGTGCGAAATACGATCCCGAAGGTGAATACATTAGACAATGGCTT<br>CCAGAACTTGCCAGGTTGCCCACGGAATGGATTCACCACCCATGGGACGCCCCTCTTACGGTTTTGAAGGCGA<br>GTGGTGTGGAGCTCGGTACCAATTACGCAAAACCGATTGTTGACATTGATACCGCGCGGGAGTTGCTGGCTAA<br>AGCCATTTCACGAACCCGGAAAGCCCAAATCATGATTGGTGCTGCACCCGGTGAATTCTCAGGCTCTGGATCC<br>CCCGGGTAA |
| >pOVC1_AtCIB_X | ACCATGGCCACCGGTAATGGAGCTATAGGAGGTGACCTTTTGCTCAATTTTCCTGACATGTCGGTCCTAGAGC<br>GCCAAAGGGCTCACCTCAAGTACCTCAATCCCACCTTTGATTCTCCTCTCGCCGGCTTCTTTGCCGATTCTTC<br>AATGATTACCGGCGGCGAGATGGACAGCTATCTTTCGACTGCCGGTTTGAATCTTCCGATGATGTACGGTGAG<br>ACGACGGTGGAAGGTGATTCAAGACTCTCAATTTCGCCGGAAACGACGCTTGGGACTGGAAATTTCAAGGCAG<br>CGAAGTTTGATACAGAGACTAAGGATTGTAATGAGGCGGCGAAGAAGATGACGATGAACAGAGATGACCTAGT<br>AGAAGAAGGAGAAGAAGAGAAGTCGAAAATAACAGAGCAAAACAATGGGAGCACAAAAAGCATCAAGAAGATG<br>AAACACAAAGCCAAGAAAGAAGAGAACAATTTCTCTAATGATTCATCTAAAGTGACGAAGGAATTGGAGAAAA<br>CGGATTATATTCATGTACCGGTGAATTCTCAGGCTCTGGATCCCCCGGGTAA |
| >pOVC1_AtPIF6_X | ACCATGGCCACCGGTATGATGTTCTTACCAACCGATTATTGTTGCAGGTTAAGCGATCAAGAGTATATGGAGC<br>TTGTGTTTGAGAATGGCCAGATTCTTGCAAAGGGCCAAAGATCCAACGTTTCTCTGCATAATCAACGTACCAA<br>ATCGATCATGGATTTGTATGAGGCAGAGTATAACGAGGATTTCATGAAGAGTATCATCCATGGTGGTGGTGGT<br>GCCATCACAAATCTCGGGGACACGCAGGTTGTTCCACAAAGTCATGTTGCTGCTGCCCATGAAACAAACATGT<br>TGGAAAGCAATAAACATGTTGACCCCGGTGAATTCTCAGGCTCTGGATCCCCCGGGTAA |
| >pOVC1_AtPHYB-S_X | ACCGCCACCGGTATGGTTTCCGGAGTCGGGGGTAGTGGCGGTGGCCGTGGCGGTGGCCGTGGCGGAGAAGAAG<br>AACCGTCGTCAAGTCACACTCCTAATAACCGAAGAGGAGGAGAACAAGCTCAATCGTCGGGAACGAAATCTCT<br>CAGACCAAGAAGCAACACTGAATCAATGAGCAAAGCAATTCAACAGTACACCGTCGACGCAAGACTCCACGCC<br>GTTTTCGAACAATCCGGCGAATCAGGGAAATCATTCGACTACTCACAATCACTCAAAACGACGACGTACGGTT<br>CCTCTGTACCTGAGCAACAGATCACAGCTTATCTCTCTCGAATCCAGCGAGGTGGTTACATTCAGCCTTTCGG<br>ATGTATGATCGCCGTCGATGAATCCAGTTTCCGGATCATCGGTTACAGTGAAAACGCCAGAGAAATGTTAGGG<br>ATTATGCCTCAATCTGTTCCTACTCTTGAGAAACCTGAGATTCTAGCTATGGGAACTGATGTGAGATCTTTGT<br>TCACTTCTTCGAGCTCGATTCTACTCGAGCGTGCTTTCGTTGCTCGAGAGATTACCTTGTTAAATCCGGTTTG<br>GATCCATTCCAAGAATACTGGTAAACCGTTTTACGCCATTCTTCATAGGATTGATGTTGGTGTTGTTATTGAT<br>TTAGAGCCAGCTAGAACTGAAGATCCTGCGCTTTCTATTGCTGGTGCTGTTCAATCGCAGAAACTCGCGGTTC<br>GTGCGATTTCTCAGTTACAGGCTCTTCCTGGTGGAGATATTAAGCTTTTGTGTGACACTGTCGTGGAAAGTGT<br>GAGGGACTTGACTGGTTATGATCGTGTTATGGTTTATAAGTTTCATGAAGATGAGCATGGAGAAGTTGTAGCT<br>GAGAGTAAACGAGATGATTTAGAGCCTTATATTGGACTGCATTATCCTGCTACTGATATTCCTCAAGCGTCAA<br>GGTTCTTGTTTAAGCAGAACCGTGTCCGAATGATAGTAGATTGCAATGCCACACCTGTTCTTGTGGTCCAGGA<br>CGATAGGCTAACTCAGTCTATGTGCTTGGTTGGTTCTACTCTTAGGGCTCCTCATGGTTGTCACTCTCAGTAT<br>ATGGCTAACATGGGATCTATTGCGTCTTTAGCAATGGCGGTTATAATCAATGGAAATGAAGATGATGGGAGCA<br>ATGTAGCTAGTGGAAGAAGCTCGATGAGGCTTTGGGGTTTGGTTGTTTGCCATCACACTTCTTCTCGCTGCAT<br>ACCGTTTCCGCTAAGGTATGCTTGTGAGTTTTTGATGCAGGCTTTCGGTTTACAGTTAAACATGGAATTGCAG<br>TTAGCTTTGCAAATGTCAGAGAAACGCGTTTTGAGAACGCAGACACTGTTATGTGATATGCTTCTGCGTGACT<br>CGCCTGCTGGAATTGTTACACAGAGTCCCAGTATCATGGACTTAGTGAAATGTGACGGTGCAGCATTTCTTTA<br>CCACGGGAAGTATTACCCGTTGGGTGTTGCTCCTAGTGAAGTTCAGATAAAAGATGTTGTGGAGTGGTTGCTT |

| | |
|---|---|
| | GCGAATCATGCGGATTCAACCGGATTAAGCACTGATAGTTTAGGCGATGCGGGGTATCCCGGTGCAGCTGCGT<br>TAGGGGATGCTGTGTGCGGTATGGCAGTTGCATATATCACAAAAAGAGACTTTCTTTTTTGGTTTCGATCTCA<br>CACTGCGAAAGAAATCAAATGGGGAGGCGCTAAGCATCATCCGGAGGATAAAGATGATGGGCAACGAATGCAT<br>CCTCGTTCGTCCTTTCAGGCTTTTCTTGAAGTTGTTAAGAGCCGGAGTCAGCCATGGGAAACTGCGGAAATGG<br>ATGCGATTCACTCGCTCCAGCTTATTCTGAGAGACTCTTTTAAAGAATCTGAGGCGGCTATGAACTCTAAAGT<br>TGTGGATGGTGTGGTTCAGCCATGTAGGGATATGGCGGGGGAACAGGGGATTGATGAGTTAGGTGCAGTTGCA<br>AGAGAGATGGTTAGGCTCATTGAGACTGCAACTGTTCCTATATTCGCTGTGGATGCCGGAGGCTGCATCAATG<br>GATGGAACGCTAAGATTGCAGAGTTGACAGGTCTCTCAGTTGAAGAAGCTATGGGGAAGTCTCTGGTTTCTGA<br>TTTAATATACAAAGAGAATGAAGCAACTGTCAATAAGCTTCTTTCTCGTGCTTTGAGAGGGGACGAGGAAAAG<br>AATGTGGAGGTTAAGCTGAAAACTTTCAGCCCCGAACTACAAGGGAAAGCAGTTTTTGTGGTTGTGAATGCTT<br>GTTCCAGCAAGGACTACTTGAACAACATTGTCGGCGTTTGTTTTGTTGGACAAGACGTTACTAGTCAGAAAAT<br>CGTAATGGATAAGTTCATCAACATACAAGGAGATTACAAGGCTATTGTACATAGCCCAAACCCTCTAATCCCG<br>CCAATTTTTGCTGCTGACGAGAACACGTGCTGCCTGGAATGGAACATGGCGATGGAAAAGCTTACGGTTGGT<br>CTCGCAGTGAAGTGATTGGGAAAATGATTGTCGGGGAAGTGTTTGGGAGCTGTTGCATGCTAAAGGGTCCTGA<br>TGCTTTAACCAAGTTCATGATTGTATTGCATAATGCGATTGGTGGCCAAGATACGGATAAGTTCCCTTTCCCA<br>TTCTTTGACCGCAATGGGAAGTTTGTTCAGGCTCTATTGACTGCAAACAAGCGGGTTAGCCTCGAGGGAAAGG<br>TTATTGGGGCTTTCTGTTTCTTGCAAATCCCGAGCCCGGTGAATTCTCAGGCTCTGGATCCCCCGGGTAA |
| >pOVC1_ScPH1-S_X | ACCATGGCCACCGGGGCAACTACTGTTCAACTGTCTGATCAATCTCTGCGTCAACTGGAAACTCTGGCTATCC<br>ACACCGCGCATCTGATCCAGCCGCACGGTCTGGTAGTCGTCCTGCAAGAACCGGACCTGACCATCAGCCAGAT<br>CTCTGCGAACTGTACCGGCATCCTGGGCCGTAGCCCGGAAGATCTGCTGGGTCGTACTCTGGGCGAGGTATTC<br>GATTCTTTTCAGATTGATCCGATCCAGTCTCGTCTGACCGCAGGTCAGATTTCCAGCCTGAACCCGTCCAAGC<br>TGTGGGCGCGTGTTATGGGTGACGACTTTGTTATTTCGACGGCGTATTTCATCGTAACTCTGATGGCCTGCT<br>GGTTTGCGAGCTGGAGCCGGCCTACACTAGCGACAACCTGCCTTTCCTGGGTTTCTACCATATGGCAAACGCG<br>GCACTGAACCGTCTGCGTCAGCAAGCTAACCTGCGCGACTTCTACGACGTTATCGTTGAGGAAGTGCGCCGCA<br>TGACGGGTTTCGACCGCGTCATGCTGTACCGTTTTGATGAAAACAACCACGGTGACGTAATCGCGGAGGATAA<br>GCGTGACGACATGGAGCCGTATCTGGGTCTGCACTACCCGGAAAGCGACATTCCTCAGCCGGCACGTCGCCTG<br>TTCATTCACAACCCGATCCGTGTTATTCCGGACGTTTACGGCGTTGCTGTTCCGCTGACTCCGGCCGTTAATC<br>CGTCTACTAACCGTGCAGTTGACCTGACCGAATCCATCCTGCGTTCCGCATACCATTGCCACCTGACCTATCT<br>GAAGAACATGGGCGTTGGTGCTAGCCTGACGATCTCTCTGATTAAAGATGGTCACCTGTGGGGTCTGATCGCT<br>TGCCATCACCAGACCCCGAAAGTAATCCCTTTCGAACTGCGTAAAGCCTGCGAATTCTTCGGTCGTGTGGTGT<br>TCTCTAATATCTCCGCGCAAGAAGACACCGAGACTTTTGACTACCGCGTACAGCTGGCGGAGCATGAAGCGGT<br>TCTGCTGGACAAAATGACCACCGCGGCAGACTTCGTGGAGGGCCTGACTAACCACCCAGACCGTCTGCTGGGC<br>CTGACCGGCAGCCAAGGCGCTGCGATTTGTTTCGGCGAGAAACTGATTCTGGTGGGCGAAACCCCAGACGAAA<br>AGGCGGTGCAATACCTGCTGCAATGGCTGGAGAATCGCGAAGTGCAGGACGTTTTCTTCACTAGCTCTCTGTC<br>TCAGATCTATCCGGATGCGGTTAACTTCAAAAGCGTGGCGTCCGGCCTGCTGGCTATCCCGATCGCCCGTCAT<br>AACTTTCTGCTGTGGTTCCGCCCGGAGGTTCTGCAGACCGTTAATTGGGGTGGTGATCCGAATCACGCATACG<br>AAGCAACCCAAGAAGATGGTAAGATCGAACTGCATCCGCGTCAGTCCTTCGATCTGTGGAAAGAAATTGTTCG<br>CCTGCAGAGCCTGCCGTGGCAGAGCGTTGAGATCCAGTCTGCCCTGGCTCTGAAGAAAGCAATCGTGAACCTG<br>ATTCTGCGCCAAGCTGAAGAACCCGGTGAATTCTCAGGCTCTGGATCCCCCGGGTAA |
| >pOVC1_X_VfAU1-LOV | ACCATGGCCACCGGTGAATTCTCAGGCTCTGGATCCCCCGGTCCTGACTACAGTCTCGTGAAGGCTCTGCAAA<br>TGGCACAACAGAATTTTGTCATTACAGACGCCTCCCTCCCAGACAACCCTATCGTCTACGCCAGTAGAGGGTT<br>TCTGACACTGACAGGCTATTCTCTCGACCAGATCCTGGGCAGGAACTGCAGGTTTCTGCAAGGGCCAGAAACA<br>GACCCAAGAGCTGTGGATAAGATCAGGAATGCCATCACCAAAGGCGTTGATACCAGTGTCTGTCTGCTGAATT<br>ATAGACAGGATGGCACAACCTTCTGGAATCTCTTCTTCGTGGCTGGACTCAGAGATTCTAAGGGCAATATTGT<br>CAACTACGTCGGAGTGCAGTCAAAGGTGAGCGAAGATTATGCCAAGCTGCTGGTCAACGAGCAGAACATTGAG<br>TACAAAGGTGTGCGCACCAGTAACATGCTGCGCAGAAAGCCCGGGTAA |
| >pOVC1_X_CrPH-LOV | ACCATGGCCACCGGTGAATTCTCAGGCTCTGGATCCCCCGGTGCAGGACTCAGACATACATTTGTGGTGGCTG<br>ATGCAACACTCCCTGATTGCCCACTGGTCTATGCAAGTGAGGGCTTCTACGCAATGACCGGATATGGACCTGA<br>CGAAGTGCTGGGTCACAACTGTAGGTTTCTGCAGGGTGAGGGAACTGACCCCAAGGAAGTGCAGAAAATTCGC<br>GACGCCATCAAGAAGGGTGAGGCTTGTAGTGTGCGCCTCCTGAACTATCGGAAGGACGGCACTCCCTTCTGGA<br>ACCTGCTGACAGTCACCCCAATTAAAACCCCTGATGGCCGCGTGTCCAAGTTTGTCGGCGTGCAGGTGGATGT<br>TACCTCCAAGACTGAAGGGAAAGCCCTGGCCCCCGGGTAA |
| >pOVC1_X_NcVVD-LOV | ACCATGGCCACCGGTGAATTCTCAGGCTCTGGATCCCCCGGGCACACTCTCTACGCCCCAGGCGGGTACGATA<br>TTATGGGCTGGCTGATCCAGATCATGAACAGGCCCAATCCCCAGGTCGAGCTGGGACCCGTGGATACTTCATG<br>TGCACTGATACTGTGCGACCTGAAGCAGAAGGATACACCTATAGTTTACGCTTCAGAAGCCTTTCTGTACATG<br>ACAGGGTATTCTAACGCCGAGGTGCTGGGGAGGAACTGTAGGTTCCTCCAGAGTCCCGATGGTATGGTGAAAC<br>CTAAGAGTACTCGCAAATATGTGGATAGCAATACTATTAACACCATGAGGAAAGCCATCGACAGAAACGCAGA<br>AGTTCAGGTGGAAGTGGTGAACTTTAAGAAGAACGGCCAGCGGTTCGTGAACTTTCTCACAATGATTCCAGTG<br>CGGGACGAAACCGGGGAGTACCGGTACAGCATGGGTTTTCAGTGCGAAACCGAACCCGGGTAA |
| >pOVC1_X_RsLP-LOV | ACCATGGCCACCGGTGAATTCTCAGGCTCTGGATCCCCCGGTGCCATGGATCAGAAGCAGTTTGAGAAGATTA<br>GAGCTGTGTTTGACAGGTCAGGGGTCGCACTGACCCTCGTTGACATGTCCCTGCCAGAGCAACCCCTGGTGCT<br>CGCCAACCCTCCATTTCTGAGAATGACTGGCTATACTGAGGGCCAGATCCTGGGATTCAACTGCAGATTTCTC<br>CAGAGAGGCGACGAAAATGCTCAGGCACGGGCTGACATCAGAGATGCCCTCAAGCTCGGAAGGGAGCTCCAGG<br>TGGTCCTCCGCAATTACAGAGCCAACGATGAACCATTTGACAATCTGCTGTTCCTGCACCCTGTCGGTGGCAG<br>ACCCGACGCTCCTGACTACTTCCTCGGTTCTCAGTTCGAGCTGGGTAGAAGCGGAAATAGCGAAGAGGCAGCC<br>GCAGCTGGACACGCAGGGGCACTGACTGGGGAGCTCGCCAGAATAGGAACTGTGGCTGCTCGGCTCGAAATGG<br>ACAGTCGGAGACATCTGGCACAAGCTGCTGCAGCCCTGGTGAGGGCCTGGGAAAGAAGGGGTCCCGGGTAA |

| | |
|---|---|
| >pOVC1_X_AsPT1-LOV2-pep | ACCATGGCCACCGGTGAATTCTCAGGCTCTGGATCCCCCGGGTTGGCTGCTGCACTTGAACGTATTGAGAAGA<br>ACTTTGTCATTACTGACCCAAGATTGCCAGATAATCCCATTATATTCGCGTCCGATAGTTTCTTGCAGTTGAC<br>AGAATATAGCCGTGAAGAAATTTTGGGAAGAAACTGCAGGTTTCTACAAGGTCCTGAAACTGATCGCGCGACA<br>GTGAGAAAAATTAGAGATGCCATAGATAACCAAACAGAGGTCACTGTTCAGCTGATTAATTATACAAAGAGTG<br>GTAAAAAGTTCTGGAACCTCTTTCACTTGCAGCCTATGCGAGATCAGAAGGGAGATGTCCAGTACTTTATTGG<br>GGTTCAGTTGGATGGAACTGAGCATGTCCGAGATGCTGCCGAGAGAGAGGCAGTCATGCTGGCGAAGAAAACT<br>GCAGAAGAGATTGATAAAGCGGTGGATACCTGGGTGCCCGGGTAA |
| >pOVC1_X_HsPDZ1b | ACCATGGCCACCGGTGAATTCTCAGGCTCTGGATCCCCCGGGCCAGAACTTGGATTTAGCATATCAGGTGGTG<br>TCGGGGGTAGAGGAAACCCATTCAGACCTGATGATGATGGTATATTTGTAACAAGGGTACAACCTGAAGGACC<br>AGCATCAAAATTACTGCAGCCAGGTGATAAAATTATTCAGGCTAATGGCTACAGTTTTATAAATATTGAACAT<br>GGACAAGCAGTGTCCTTGCTAAAAACTTTCCAGAATACAGTTGAACTCATCATTGTACGAGAAGTTGGTAACG<br>GTGCAAAACAAGAGATTCGAGTGAGGGTTGAAAAAGACGGTGGTTCTGGAGGCGTTTCTTCTGTTCCGACCAA<br>TCTGGAAGTTGTTGCTGCGACCCCGACTAGCCTGCTGATCAGCTGGGATGCGTATAGGGAGCTTCCGGTTAGT<br>TATTACCGTATCACGTACGGTGAAACAGGTGGTAACTCCCCGGTTCAGGAGTTCACTGTACCTGGTTCCAAGT<br>CTACTGCTACCATCAGCGGCCTGAAACCGGGTGTCGACTATACCATCACTGTATACGCACATTACAACTACCA<br>TTACTACTCTAGCCCAATCTCGATTAACTACCGTACCTCTAGACCCGGGTAA |
| >pOVC1_X_AsPT1-LOV2-EcSsra | ACCATGGCCACCGGTGAATTCTCAGGCTCTGGATCCCCCGGGCTGGCAACCACACTGGAACGGATCGAGAAAA<br>ATTTCGTGATTACTGATCCGAGACTGCCTGACAACCCAATCATTTTTGCGAGCGATTCCTTCCTGCAGCTGAC<br>AGAATATTCTCGGGAAGAGATCCTGGGGCGCAATTGCCGTTTTCTGCAGGGACCCGAGACAGACCGTGCCACT<br>GTTCGGAAAATCAGAGATGCTATTGACAACCAGACTGAAGTGACCGTTCAGCTGATCAATTATACCAAGAGCG<br>GCAAGAAGTTCTGGAACGTGTTCCACCTGCAGCCGATGCGCGATTATAAGGGCGACGTCCAGTACTTCATTGG<br>CGTGCAGCTGGATGGCACCGAACGTCTTCATGGCGCCGCTGAGCGTGAGGCGGTCTGCCTGATCAAAAAGACA<br>GCCTTTCAGATTGCTGAGGCAGCGAACGACGAAAATTACTTTGGACCCGGGTAA |
| >pOVC1_X_EcSSPB-micro | ACCATGGCCACCGGTGAATTCTCAGGCTCTGGATCCCCCGGGAGCTCCCCGAAACGCCCTAAGCTGCTGCGTG<br>AATATTACGATTGGCTGGTTGATAACAGCTTTACCCCATATCTGGTGGTGGATGCCACATACCTGGGCGTGAA<br>CGTGCCCGTGGAGTATGTGAAAGACGGTCAGATCGTGCTGAATCTGTCTGCAAGTGCGACCGGCAACCTGCAA<br>CTGACAAATGATTTTATCCAGTTCAACGCCCAGTTTAAGGGCGTGTCTCGTGAACTGTATATCCCGATGGGTG<br>CCGCTCTGGCCATTTACGCTCGCGAGAACGGCGATGGTGTGATGTTCGAACCAGAAGAAATCTATGACGAGCT<br>GAATATTGGTCCCGGGTAA |
| >pOVC1_X_EcSSPB-nano | ACCATGGCCACCGGTGAATTCTCAGGCTCTGGATCCCCCGGGAGCTCCCCGAAACGCCCTAAGCTGCTGCGTG<br>AATATTACGATTGGCTGGTTGATAACAGCTTTACCCCATATCTGGTGGTGGATGCCACATACCTGGGCGTGAA<br>CGTGCCCGTGGAGTATGTGAAAGACGGTCAGATCGTGCTGAATCTGTCTGCAAGTGCGACCGGCAACCTGCAA<br>CTGACAAATGATTTTATCCAGTTCAACGCCCGCTTTAAGGGCGTGTCTCGTGAACTGTATATCCCGATGGGTG<br>CCGCTCTGGCCATTTACGCTCGCGAGAACGGCGATGGTGTGATGTTCGAACCAGAAGAAATCTATGACGAGCT<br>GAATATTGGTCCCGGGTAA |
| pOVC1_X_MxCarH-CBD | >ACCATGGCCACCGGTGAATTCTCAGGCTCTGGATCCCCCGGTCCACACGCAGAGACTTGGAGGGAATCTATG<br>CTTGCCGCCACACAAGCCTACGACCAGCCTAGAGTATCAGATGTACTGGATGAGGTCCTTGCGGCTCTGCCCC<br>CTCTGAAGGCCTTCGATGAAGTGCTGGCCCCTTTGCTGTGCGATGTCGGAGAGCGGTGGGAGAGCGGAACCCT<br>GACAGTTGCGCAGGAACATCTGGTCTCACAGATGGTGCGCGCCCGGCTGGTGAGTCTGCTGCACGCGGCACCA<br>TTGGGACGCCACAGACATGGCGTTCTCGCCTGTTTCCCAGAGGAGGAGCATGAGATGGGCTTGCTTGGTGCCG<br>CCTTGAGACTCCGCCATCTCGGCGTTAGAGTAACCCTGCTCGGCCAGCGAGTGCCAGCCGAGGACCTCGGGCG<br>AGCAGTGTTGGCCCTGCGCCCGGACTTCGTGGCCTGTCAACAGTTGCAAGCAGGAGCGCAGAGGACTTCGAG<br>GATACCTTGACCCGACTCCGCCAGGCCCTGCCAAGGGGCCTCCCTGTATGGGTGGGCGGGGCAGCCGCAAGGT<br>CTCATCAGGCCGTGTGCGAGCGCCTGGCAGTCCATGTTTTTCAGGGCGAAGAAGATTGGGATAGACTTGCCGG<br>AACACCCGGGTAA |
| >pOVC1_X_TtCarH-CBD | ACCATGGCCACCGGTGAATTCTCAGGCTCTGGATCCCCCGGTCCCGAGGACCTCGGCACCGGACTCCTCGAAG<br>CACTCTTGAGAGGAGATTTGGCCGGCGCCGAGGCTCTCTTTCGACGGGGGCTTAGGTTTTGGGACCCGAGGG<br>TATTCTGGAGCACCTGCTCCTGCCTGTGCTTCGGGAGGTGGGAGAAGCCTGGCATCGCGGCGAGATCGGTGTG<br>GCCGAGGAGCATCTGGCATCCACATTTCTGCGCGCGAGACTGCAGGAGCTGCTCGACCTCGCCGGGTTCCCAC<br>CTGGCCCCCCCGTGCTGGTAACCACGCCACCAGGGGAGAGGCACGAGATCGGCGCAATGTTGGCTGCCTATCA<br>CCTGCGAAGGAAGGGCGTGCCAGCGCTGTACTTGGGACCAGACACCCCCCTCCCCGATCTCAGAGCACTTGCG<br>AGACGGCTCGGAGCCGGAGCGGTGGTTCTGTCAGCTTTGCTTTCCGAGCCTCTCAGGGCGTTGCCAGACGGCG<br>CACTGAAAGACTTGGCACCTCGGGTGTTCCTGGGAGGGCAAGGAGCCGGCCCTGAAGAGGCCCGACGGCTCGG<br>GGCCGAGTACATGGAAGATCTGAAGGGATTGGCCGAAGCACTGTGGCTTCCAAGAGGACCAGAGAAAGAAGCA<br>ATCCCCGGGTAA |
| >pOVC1_X_AtCRY2-PHR | ACCATGGCCACCGGTGAATTCTCAGGCTCTGGATCCCCCGGTAAGATGGACAAGAAAACCATAGTCTGGTTTC<br>GCCGCGATTTGAGGATAGAGGATAACCCTGCGTTGGCCGCAGCGGCCCACGAAGGCAGCGTGTTCCCCGTGTT<br>CATATGGTGCCCAGAGGAGGAAGGCCAGTTCTACCCAGGTCGCGCTAGTCGCTGGTGGATGAAACAGTCTCTC<br>GCACATCTCTCCCAATCTTTGAAAGCTCTCGGGTCTGACCTTACGCTGATTAAGACCCACAATACTATTAGTG<br>CAATCTTGGACTGCATCCGCGTTACCGGCGCGACCAAAGTGGTGTTTAATCATCTGTATGACCCGGTAAGTCT<br>GGTACGCGACCATACAGTTAAGGAGAAGTTGGTTGAAAGGGGAATTTCAGTACAGAGTTATAATGGAGACCTC<br>CTCTACGAACCTTGGGAAATATACTGTGAGAAGGGAAAGCCATTTACATCATTTAACTCATACTGGAAAAAGT<br>GTCTTGACATGAGCATAGAGTCTGTCATGTTGCCCCCGCCGTGGCGGCTTATGCCCATCACCGCGGCCGCCGA<br>AGCTATCTGGGCGTGTAGTATTGAAGAACTTGGCTTGGAGAACGAAGCAGAAAAACCTAGCAATGCACTGTTG<br>ACGCGCGCCTGGTCCCCTGGGTGGAGTAACGCAGATAAATTGCTTAACGAGTTCATCGAAAAGCAACTTATCG<br>ACTACGCTAAGAATAGCAAAAAAGTGGTGGGCAATTCAACCTCCCTCCTGTCTCCGTACTTGCATTTCGGCGA<br>GATTAGCGTGCGGCACGTCTTCCAATGCGCCCGAATGAAACAAATAATCTGGGCGCGGGATAAAAACAGTGAG |

| | |
|---|---|
| | GGTGAAGAAAGCGCAGACTTGTTCCTCCGAGGAATCGGTCTGCGAGAATATAGTCGCTACATTTGTTTCAATT<br>TTCCCTTTACGCATGAGCAGAGCCTCCTTAGTCACTTGCGATTCTTTCCTTGGGACGCAGATGTTGACAAATT<br>TAAAGCATGGCGGCAAGGTAGAACAGGCTACCCATTGGTAGATGCGGGTATGCGAGAACTCTGGGCCACGGGG<br>TGGATGCATAACCGAATCAGGGTAATAGTAAGTAGTTTCGCAGTTAAGTTTCTTTTGCTTCCATGGAAGTGGG<br>GGATGAAGTATTTCTGGGACACTTTGCTCGATGCGGATCTTGAATGCGATATATTGGGTTGGCAATATATTTC<br>CGGGTCAATCCCTGACGGCCATGAGCTGGACAGATTGGACAATCCTGCGCTCCAGGGTGCGAAATACGATCCC<br>GAAGGTGAATACATTAGACAATGGCTTCCAGAACTTGCCAGGTTGCCCACGGAATGGATTCACCACCCATGGG<br>ACGCCCCTCTTACGGTTTTGAAGGCGAGTGGTGTGGAGCTCGGTACCAATTACGCAAAACCGATTGTTGACAT<br>TGATACCGCGCGGGAGTTGCTGGCTAAAGCCATTTCACGAACCCGGAAAGCCCAAATCATGATTGGTGCTGCA<br>ACCGGGTAA |
| >pOVC1_X_AtPIF6 | ACCATGGCCACCGGTGAATTCTCAGGCTCTGGATCCCCCGGTATGATGTTCTTACCAACCGATTATTGTTGCA<br>GGTTAAGCGATCAAGAGTATATGGAGCTTGTGTTTGAGAATGGCCAGATTCTTGCAAAGGGCCAAAGATCCAA<br>CGTTTCTCTGCATAATCAACGTACCAAATCGATCATGGATTTGTATGAGGCAGAGTATAACGAGGATTTCATG<br>AAGAGTATCATCCATGGTGGTGGTGGTGCCATCACAAATCTCGGGGACACGCAGGTTGTTCCACAAAGTCATG<br>TTGCTGCTGCCCATGAAACAAACATGTTGGAAAGCAATAAACATGTTGACCCCGGGTAA |
| >pOVC1_X_ScPH1-S | ACCATGGCCACCGGTGAATTCTCAGGCTCTGGATCCCCCGGGGCAACTACTGTTCAACTGTCTGATCAATCTC<br>TGCGTCAACTGGAAACTCTGGCTATCCACACCGCGCATCTGATCCAGCCGCACGGTCTGGTAGTCGTCCTGCA<br>AGAACCGGACCTGACCATCAGCCAGATCTCTGCGAACTGTACCGGCATCCTGGGCCGTAGCCCGGAAGATCTG<br>CTGGGTCGTACTCTGGGCGAGGTATTCGATTCTTTTCAGATTGATCCGATCCAGTCTCGTCTGACCGCAGGTC<br>AGATTTCCAGCCTGAACCCGTCCAAGCTGTGGGCGCGTGTTATGGGTGACGACTTTGTTATTTTCGACGGCGT<br>ATTTCATCGTAACTCTGATGGCCTGCTGGTTTGCGAGCTGGAGCCGGCCTACACTAGCGACAACCTGCCTTTC<br>CTGGGTTTCTACCATATGGCAAACGCGGCACTGAACCGTCTGCGTCAGCAAGCTAACCTGCGCGACTTCTACG<br>ACGTTATCGTTGAGGAAGTGCGCCGCATGACGGGTTTCGACCGCGTCATGCTGTACCGTTTTGATGAAAACAA<br>CCACGGTGACGTAATCGCGGAGGATAAGCGTGACGACATGGAGCCGTATCTGGGTCTGCACTACCCGGAAAGC<br>GACATTCCTCAGCCGGCACGTCGCCTGTTCATTCACAACCCGATCCGTGTTATTCCGGACGTTTACGGCGTTG<br>CTGTTCCGCTGACTCCGGCCGTTAATCCGTCTACTAACCGTGCAGTTGACCTGACCGAATCCATCCTGCGTTC<br>CGCATACCATTGCCACCTGACCTATCTGAAGAACATGGGCGTTGGTGCTAGCCTGACGATCTCTCTGATTAAA<br>GATGGTCACCTGTGGGGTCTGATCGCTTGCCATCACCAGACCCCGAAAGTAATCCCTTTCGAACTGCGTAAAG<br>CCTGCGAATTCTTCGGTCGTGTGGTGTTCTCTAATATCTCCGCGCAAGAAGACACCGAGACTTTTGACTACCG<br>CGTACAGCTGGCGGAGCATGAAGCGGTTCTGCTGGACAAAATGACCACCGCGGCAGACTTCGTGGAGGGCCTG<br>ACTAACCACCCAGACCGTCTGCTGGGCCTGACCGGCAGCCAAGGCGCTGCGATTTGTTTCGGCGAGAAACTGA<br>TTCTGGTGGGCGAAACCCCAGACGAAAAGGCGGTGCAATACCTGCTGCAATGGCTGGAGAATCGCGAAGTGCA<br>GGACGTTTTCTTCACTAGCTCTCTGTCTCAGATCTATCCGGATGCGGTTAACTTCAAAAGCGTGGCGTCCGGC<br>CTGCTGGCTATCCCGATCGCCCGTCATAACTTTCTGCTGTGGTTCCGCCCGGAGGTTCTGCAGACCGTTAATT<br>GGGGTGGTGATCCGAATCACGCATACGAAGCAACCCAAGAAGATGGTAAGATCGAACTGCATCCGCGTCAGTC<br>CTTCGATCTGTGGAAAGAAATTGTTCGCCTGCAGAGCCTGCCGTGGCAGAGCGTTGAGATCCAGTCTGCCCTG<br>GCTCTGAAGAAAGCAATCGTGAACCTGATTCTGCGCCAAGCTGAAGAACCCGGGTAA |
| >pOVC1_X_AtCIB | ACCATGGCCACCGGTGAATTCTCAGGCTCTGGATCCCCCGGTAATGGAGCTATAGGAGGTGACCTTTTGCTCA<br>ATTTTCCTGACATGTCGGTCCTAGAGCGCCAAAGGGCTCACCTCAAGTACCTCAATCCCACCTTTGATTCTCC<br>TCTCGCCGGCTTCTTTGCCGATTCTTCAATGATTACCGGCGGCGAGATGGACAGCTATCTTTCGACTGCCGGT<br>TTGAATCTTCCGATGATGTACGGTGAGACGACGGTGGAAGGTGATTCAAGACTCTCAATTTCGCCGGAAACGA<br>CGCTTGGGACTGGAAATTTCAAGGCAGCGAAGTTTGATACAGAGACTAAGGATTGTAATGAGGCGGCGAAGAA<br>GATGACGATGAACAGAGATGACCTAGTAGAAGAAGGAGAAGAAGAGAAGTCGAAAATAACAGAGCAAAACAAT<br>GGGAGCACAAAAAGCATCAAGAAGATGAAACACAAAGCCAAGAAAGAAGAGAACAATTTCTCTAATGATTCAT<br>CTAAAGTGACGAAGGAATTGGAGAAAACGGATTATATTCATGTCCCGGGTAA |

**Supplementary Table 6.** Nucleotide sequences for Opto-casp9 constructs. Green: site *A*, blue: site *B*, red: site *C*. The genes are available through Addgene.org.

| Name | Sequence |
|---|---|
| >pOVC1_VfAU1-LOV_casp9 | ACCATGGCCACCGGTCCTGACTACAGTCTCGTGAAGGCTCTGCAAATGGCACAACAGAATTTTGTCATTACAGACGCCTC<br>CCTCCCAGACAACCCTATCGTCTACGCCAGTAGAGGGTTTCTGACACTGACAGGCTATTCTCTCGACCAGATCCTGGGCA<br>GGAACTGCAGGTTTCTGCAAGGGCCAGAAACAGACCCAAGAGCTGTGGATAAGATCAGGAATGCCATCACCAAAGGCGTT<br>GATACCAGTGTCTGTCTGCTGAATTATAGACAGGATGGCACAACCTTCTGGAATCTCTTCTTCGTGGCTGGACTCAGAGA<br>TTCTAAGGGCAATATTGTCAACTACGTCGGAGTGCAGTCAAAGGTGAGCGAAGATTATGCCAAGCTGCTGGTCAACGAGC<br>AGAACATTGAGTACAAAGGTGTGCGCACCAGTAACATGCTGCGCAGAAAGCCCGGTGAATTCTCAGGCTCTGGATCCCCC<br>GGGGGATTTGGTGATGTCGGTGCTCTTGAGAGTTTGAGGGGAAATGCAGATTTGGCTTACATCCTGAGCATGGAGCCCTG<br>TGGCCACTGCCTCATTATCAACAATGTGAACTTCTGCCGTGAGTCCGGGCTCCGCACCCGCACTGGCTCCAACATCGACT<br>GTGAGAAGTTGCGGCGTCGCTTCTCCTCGCTGCATTTCATGGTGGAGGTGAAGGGCGACCTGACTGCCAAGAAAATGGTG<br>CTGGCTTTGCTGGAGCTGGCGCGGCAGGACCACGGTGCTCTGGACTGCTGCGTGGTGGTCATTCTCTCTCACGGCTGTCA<br>GGCCAGCCACCTGCAGTTCCCAGGGGCTGTCTACGGCACAGATGGATGCCCTGTGTCGGTCGAGAAGATTGTGAACATCT<br>TCAATGGGACCAGCTGCCCCAGCCTGGGAGGGAAGCCCAAGCTCTTTTTCATCCAGGCCTGTGGTGGGGAGCAGAAAGAC<br>CATGGGTTTGAGGTGGCCTCCACTTCCCCTGAAGACGAGTCCCCTGGCAGTAACCCCGAGCCAGATGCCACCCCGTTCCA<br>GGAAGGTTTGAGGACCTTCGACCAGCTGGACGCCATATCTAGTTTGCCCACACCCAGTGACATCTTTGTGTCCTACTCTA<br>CTTTCCCAGGTTTTGTTTCCTGGAGGGACCCCAAGAGTGGCTCCTGGTACGTTGAGACCCTGGACGACATCTTTGAGCAG<br>TGGGCTCACTCTGAAGACCTGCAGTCCCTCCTGCTTAGGGTCGCTAATGCTGTTTCGGTGAAAGGGATTTATAAACAGAT<br>GCCTGGTTGCTTTAATTTCCTCCGGAAAAAACTTTTCTTTAAAACATCACCCGGGTAA |
| >pOVC1_CrPH-LOV_casp9 | ACCATGGCCACCGGTGCAGGACTCAGACATACATTTGTGGTGGCTGATGCAACACTCCCTGATTGCCCACTGGTCTATGC<br>AAGTGAGGGCTTCTACGCAATGACCGGATATGGACCTGACGAAGTGCTGGGTCACAACTGTAGGTTTCTGCAGGGTGAGG<br>GAACTGACCCCAAGGAAGTGCAGAAAATTCGCGACGCCATCAAGAAGGGTGAGGCTTGTAGTGTGCGCCTCCTGAACTAT<br>CGGAAGGACGGCACTCCCTTCTGGAACCTGCTGACAGTCACCCCAATTAAAACCCCTGATGGCCGCGTGTCCAAGTTTGT<br>CGGCGTGCAGGTGGATGTTACCTCCAAGACTGAAGGGAAAGCCCTGGCCCCCGGTGAATTCTCAGGCTCTGGATCCCCCC<br>GGGGGATTTGGTGATGTCGGTGCTCTTGAGAGTTTGAGGGGAAATGCAGATTTGGCTTACATCCTGAGCATGGAGCCCTG<br>TGGCCACTGCCTCATTATCAACAATGTGAACTTCTGCCGTGAGTCCGGGCTCCGCACCCGCACTGGCTCCAACATCGACT<br>GTGAGAAGTTGCGGCGTCGCTTCTCCTCGCTGCATTTCATGGTGGAGGTGAAGGGCGACCTGACTGCCAAGAAAATGGTG<br>CTGGCTTTGCTGGAGCTGGCGCGGCAGGACCACGGTGCTCTGGACTGCTGCGTGGTGGTCATTCTCTCTCACGGCTGTCA<br>GGCCAGCCACCTGCAGTTCCCAGGGGCTGTCTACGGCACAGATGGATGCCCTGTGTCGGTCGAGAAGATTGTGAACATCT<br>TCAATGGGACCAGCTGCCCCAGCCTGGGAGGGAAGCCCAAGCTCTTTTTCATCCAGGCCTGTGGTGGGGAGCAGAAAGAC<br>CATGGGTTTGAGGTGGCCTCCACTTCCCCTGAAGACGAGTCCCCTGGCAGTAACCCCGAGCCAGATGCCACCCCGTTCCA<br>GGAAGGTTTGAGGACCTTCGACCAGCTGGACGCCATATCTAGTTTGCCCACACCCAGTGACATCTTTGTGTCCTACTCTA<br>CTTTCCCAGGTTTTGTTTCCTGGAGGGACCCCAAGAGTGGCTCCTGGTACGTTGAGACCCTGGACGACATCTTTGAGCAG<br>TGGGCTCACTCTGAAGACCTGCAGTCCCTCCTGCTTAGGGTCGCTAATGCTGTTTCGGTGAAAGGGATTTATAAACAGAT<br>GCCTGGTTGCTTTAATTTCCTCCGGAAAAAACTTTTCTTTAAAACATCACCCGGGTAA |
| >pOVC1_NcVVD-LOV_casp9 | ACCATGGCCACCGGGCACACTCTCTACGCCCCAGGCGGGTACGATATTATGGGCTGGCTGATCCAGATCATGAACAGGCC<br>CAATCCCCAGGTCGAGCTGGGACCCGTGGATACTTCATGTGCACTGATACTGTGCGACCTGAAGCAGAAGGATACACCTA<br>TAGTTTACGCTTCAGAAGCCTTTCTGTACATGACAGGGTATTCTAACGCCGAGGTGCTGGGGAGGAACTGTAGGTTCCTC<br>CAGAGTCCCGATGGTATGGTGAAACCTAAGAGTACTCGCAAATATGTGGATAGCAATACTATTAACACCATGAGGAAAGC<br>CATCGACAGAAACGCAGAAGTTCAGGTGGAAGTGGTGAACTTTAAGAAGAACGGCCAGCGGTTCGTGAACTTTCTCACAA<br>TGATTCCAGTGCGGGACGAAACCGGGGAGTACCGGTACAGCATGGGTTTTCAGTGCGAAACCGAACCCGGTGAATTCTCA<br>GGCTCTGGATCCCCCGGATTTGGTGATGTCGGTGCTCTTGAGAGTTTGAGGGGAAATGCAGATTTGGCTTACATCCTGAG<br>CATGGAGCCCTGTGGCCACTGCCTCATTATCAACAATGTGAACTTCTGCCGTGAGTCCGGGCTCCGCACCCGCACTGGCT<br>CCAACATCGACTGTGAGAAGTTGCGGCGTCGCTTCTCCTCGCTGCATTTCATGGTGGAGGTGAAGGGCGACCTGACTGCC<br>AAGAAAATGGTGCTGGCTTTGCTGGAGCTGGCGCGGCAGGACCACGGTGCTCTGGACTGCTGCGTGGTGGTCATTCTCTC<br>TCACGGCTGTCAGGCCAGCCACCTGCAGTTCCCAGGGGCTGTCTACGGCACAGATGGATGCCCTGTGTCGGTCGAGAAGA<br>TTGTGAACATCTTCAATGGGACCAGCTGCCCCAGCCTGGGAGGGAAGCCCAAGCTCTTTTTCATCCAGGCCTGTGGTGGG<br>GAGCAGAAAGACCATGGGTTTGAGGTGGCCTCCACTTCCCCTGAAGACGAGTCCCCTGGCAGTAACCCCGAGCCAGATGC<br>CACCCCGTTCCAGGAAGGTTTGAGGACCTTCGACCAGCTGGACGCCATATCTAGTTTGCCCACACCCAGTGACATCTTTG<br>TGTCCTACTCTACTTTCCCAGGTTTTGTTTCCTGGAGGGACCCCAAGAGTGGCTCCTGGTACGTTGAGACCCTGGACGAC<br>ATCTTTGAGCAGTGGGCTCACTCTGAAGACCTGCAGTCCCTCCTGCTTAGGGTCGCTAATGCTGTTTCGGTGAAAGGGAT<br>TTATAAACAGATGCCTGGTTGCTTTAATTTCCTCCGGAAAAAACTTTTCTTTAAAACATCAGGGTAA |
| >pOVC1_RsLP-LOV_casp9 | ACCATGGCCACCGGTGCCATGGATCAGAAGCAGTTTGAGAAGATTAGAGCTGTGTTTGACAGGTCAGGGGTCGCACTGAC<br>CCTCGTTGACATGTCCCTGCCAGAGCAACCCCTGGTGCTCGCCAACCCTCCATTTCTGAGAATGACTGGCTATACTGAGG<br>GCCAGATCCTGGGATTCAACTGCAGATTTCTCCAGAGAGGCGACGAAAATGCTCAGGCACGGGCTGACATCAGAGATGCC<br>CTCAAGCTCGGAAGGGAGCTCCAGGTGGTCCTCCGCAATTACAGAGCCAACGATGAACCATTTGACAATCTGCTGTTCCT<br>GCACCCTGTCGGTGGCAGACCCGACGCTCCTGACTACTTCCTCGGTTCTCAGTTCGAGCTGGGTAGAAGCGGAAATAGCG<br>AAGAGGCAGCCGCAGCTGGACACGCAGGGGCACTGACTGGGGAGCTCGCCAGAATAGGAACTGTGGCTGCTCGGCTCGAA<br>ATGGACAGTCGGAGACATCTGGCACAAGCTGCTGCAGCCCTGGTGAGGGCCTGGGAAAGAAGGGGTCCCGGTGAATTCTC<br>AGGCTCTGGATCCCCCGGGGGATTTGGTGATGTCGGTGCTCTTGAGAGTTTGAGGGGAAATGCAGATTTGGCTTACATCC<br>TGAGCATGGAGCCCTGTGGCCACTGCCTCATTATCAACAATGTGAACTTCTGCCGTGAGTCCGGGCTCCGCACCCGCACT<br>GGCTCCAACATCGACTGTGAGAAGTTGCGGCGTCGCTTCTCCTCGCTGCATTTCATGGTGGAGGTGAAGGGCGACCTGAC<br>TGCCAAGAAAATGGTGCTGGCTTTGCTGGAGCTGGCGCGGCAGGACCACGGTGCTCTGGACTGCTGCGTGGTGGTCATTC<br>TCTCTCACGGCTGTCAGGCCAGCCACCTGCAGTTCCCAGGGGCTGTCTACGGCACAGATGGATGCCCTGTGTCGGTCGAG<br>AAGATTGTGAACATCTTCAATGGGACCAGCTGCCCCAGCCTGGGAGGGAAGCCCAAGCTCTTTTTCATCCAGGCCTGTGG<br>TGGGGAGCAGAAAGACCATGGGTTTGAGGTGGCCTCCACTTCCCCTGAAGACGAGTCCCCTGGCAGTAACCCCGAGCCAG |

| | |
|---|---|
| | ATGCCACCCCGTTCCAGGAAGGTTTGAGGACCTTCGACCAGCTGGACGCCATATCTAGTTTGCCCACACCCAGTGACATC<br>TTTGTGTCCTACTCTACTTTCCCAGGTTTTGTTTCCTGGAGGGACCCCAAGAGTGGCTCCTGGTACGTTGAGACCCTGGA<br>CGACATCTTTGAGCAGTGGGCTCACTCTGAAGACCTGCAGTCCCTCCTGCTTAGGGTCGCTAATGCTGTTTCGGTGAAAG<br>GGATTTATAAACAGATGCCTGGTTGCTTTAATTTCCTCCGGAAAAAACTTTTCTTTAAAACATCACCCGGGTAA |
| >pOVC1_AtCRY2-PHR_casp9 | ACCATGGCCACCGGTAAGATGGACAAGAAAACCATAGTCTGGTTTCGCCGCGATTTGAGGATAGAGGATAACCCTGCGTT<br>GGCCGCAGCGGCCCACGAAGGCAGCGTGTTCCCCGTGTTCATATGGTGCCCAGAGGAGGAAGGCCAGTTCTACCCAGGTC<br>GCGCTAGTCGCTGGTGGATGAAACAGTCTCTCGCACATCTCTCCCAATCTTTGAAAGCTCTCGGGTCTGACCTTACGCTG<br>ATTAAGACCCACAATACTATTAGTGCAATCTTGGACTGCATCCGCGTTACCGGCGCGACCAAAGTGGTGTTTAATCATCT<br>GTATGACCCGGTAAGTCTGGTACGCGACCATACAGTTAAGGAGAAGTTGGTTGAAAGGGGAATTTCAGTACAGAGTTATA<br>ATGGAGACCTCCTCTACGAACCTTGGGAAATATACTGTGAGAAGGGAAAGCCATTTACATCATTTAACTCATACTGGAAA<br>AAGTGTCTTGACATGAGCATAGAGTCTGTCATGTTGCCCCCGCCGTGGCGGCTTATGCCCATCACCGCGGCCGCCGAAGC<br>TATCTGGGCGTGTAGTATTGAAGAACTTGGCTTGGAGAACGAAGCAGAAAAACCTAGCAATGCACTGTTGACGCGCGCCT<br>GGTCCCCTGGGTGGAGTAACGCAGATAAATTGCTTAACGAGTTCATCGAAAAGCAACTTATCGACTACGCTAAGAATAGC<br>AAAAAAGTGGTGGGCAATTCAACCTCCCTCCTGTCTCCGTACTTGCATTTCGGCGAGATTAGCGTGCGGCACGTCTTCCA<br>ATGCGCCCGAATGAAACAAATAATCTGGGCGCGGGATAAAAACAGTGAGGGTGAAGAAAGCGCAGACTTGTTCCTCCGAG<br>GAATCGGTCTGCGAGAATATAGTCGCTACATTTGTTTCAATTTTCCCTTTACGCATGAGCAGAGCCTCCTTAGTCACTTG<br>CGATTCTTTCCTTGGGACGCAGATGTTGACAAATTTAAAGCATGGCGGCAAGGTAGAACAGGCTACCCATTGGTAGATGC<br>GGGTATGCGAGAACTCTGGGCCACGGGGTGGATGCATAACCGAATCAGGGTAATAGTAAGTAGTTTCGCAGTTAAGTTTC<br>TTTTGCTTCCATGGAAGTGGGGGATGAAGTATTTCTGGGACACTTTGCTCGATGCGGATCTTGAATGCGATATATTGGGT<br>TGGCAATATATTTCCGGGTCAATCCCTGACGGCCATGAGCTGGACAGATTGGACAATCCTGCGCTCCAGGGTGCGAAATA<br>CGATCCCGAAGGTGAATACATTAGACAATGGCTTCCAGAACTTGCCAGGTTGCCCACGGAATGGATTCACCACCCATGGG<br>ACGCCCCTCTTACGGTTTTGAAGGCGAGTGGTGTGGAGCTCGGTACCAATTACGCAAAACCGATTGTTGACATTGATACC<br>GCGCGGGAGTTGCTGGCTAAAGCCATTTCACGAACCCGGAAAGCCCAAATCATGATTGGTGCTGCACCCGGTGAATTCTC<br>AGGCTCTGGATCCCCCGGATTTGGTGATGTCGGTGCTCTTGAGAGTTTGAGGGGAAATGCAGATTTGGCTTACATCCTGA<br>GCATGGAGCCCTGTGGCCACTGCCTCATTATCAACAATGTGAACTTCTGCCGTGAGTCCGGGCTCCGCACCCGCACTGGC<br>TCCAACATCGACTGTGAGAAGTTGCGGCGTCGCTTCTCCTCGCTGCATTTCATGGTGGAGGTGAAGGGCGACCTGACTGC<br>CAAGAAAATGGTGCTGGCTTTGCTGGAGCTGGCGCGGCAGGACCACGGTGCTCTGGACTGCTGCGTGGTGGTCATTCTCT<br>CTCACGGCTGTCAGGCCAGCCACCTGCAGTTCCCAGGGGCTGTCTACGGCACAGATGGATGCCCTGTGTCGGTCGAGAAG<br>ATTGTGAACATCTTCAATGGGACCAGCTGCCCCAGCCTGGGAGGGAAGCCCAAGCTCTTTTTCATCCAGGCCTGTGGTGG<br>GGAGCAGAAAGACCATGGGTTTGAGGTGGCCTCCACTTCCCCTGAAGACGAGTCCCCTGGCAGTAACCCCGAGCCAGATG<br>CCACCCCGTTCCAGGAAGGTTTGAGGACCTTCGACCAGCTGGACGCCATATCTAGTTTGCCCACACCCAGTGACATCTTT<br>GTGTCCTACTCTACTTTCCCAGGTTTTGTTTCCTGGAGGGACCCCAAGAGTGGCTCCTGGTACGTTGAGACCCTGGACGA<br>CATCTTTGAGCAGTGGGCTCACTCTGAAGACCTGCAGTCCCTCCTGCTTAGGGTCGCTAATGCTGTTTCGGTGAAAGGGA<br>TTTATAAACAGATGCCTGGTTGCTTTAATTTCCTCCGGAAAAAACTTTTCTTTAAAACATCAGGGTAA |
| >pOVC1_casp9_VfAU1-LOV | ACCATGGCCACCGGGGGATTTGGTGATGTCGGTGCTCTTGAGAGTTTGAGGGGAAATGCAGATTTGGCTTACATCCTGAG<br>CATGGAGCCCTGTGGCCACTGCCTCATTATCAACAATGTGAACTTCTGCCGTGAGTCCGGGCTCCGCACCCGCACTGGCT<br>CCAACATCGACTGTGAGAAGTTGCGGCGTCGCTTCTCCTCGCTGCATTTCATGGTGGAGGTGAAGGGCGACCTGACTGCC<br>AAGAAAATGGTGCTGGCTTTGCTGGAGCTGGCGCGGCAGGACCACGGTGCTCTGGACTGCTGCGTGGTGGTCATTCTCTC<br>TCACGGCTGTCAGGCCAGCCACCTGCAGTTCCCAGGGGCTGTCTACGGCACAGATGGATGCCCTGTGTCGGTCGAGAAGA<br>TTGTGAACATCTTCAATGGGACCAGCTGCCCCAGCCTGGGAGGGAAGCCCAAGCTCTTTTTCATCCAGGCCTGTGGTGGG<br>GAGCAGAAAGACCATGGGTTTGAGGTGGCCTCCACTTCCCCTGAAGACGAGTCCCCTGGCAGTAACCCCGAGCCAGATGC<br>CACCCCGTTCCAGGAAGGTTTGAGGACCTTCGACCAGCTGGACGCCATATCTAGTTTGCCCACACCCAGTGACATCTTTG<br>TGTCCTACTCTACTTTCCCAGGTTTTGTTTCCTGGAGGGACCCCAAGAGTGGCTCCTGGTACGTTGAGACCCTGGACGAC<br>ATCTTTGAGCAGTGGGCTCACTCTGAAGACCTGCAGTCCCTCCTGCTTAGGGTCGCTAATGCTGTTTCGGTGAAAGGGAT<br>TTATAAACAGATGCCTGGTTGCTTTAATTTCCTCCGGAAAAAACTTTTCTTTAAAACATCACCCGGTGAATTCTCAGGCT<br>CTGGATCCCCCGGTCCTGACTACAGTCTCGTGAAGGCTCTGCAAATGGCACAACAGAATTTTGTCATTACAGACGCCTCC<br>CTCCCAGACAACCCTATCGTCTACGCCAGTAGAGGGTTTCTGACACTGACAGGCTATTCTCTCGACCAGATCCTGGGCAG<br>GAACTGCAGGTTTCTGCAAGGGCCAGAAACAGACCCAAGAGCTGTGGATAAGATCAGGAATGCCATCACCAAAGGCGTTG<br>ATACCAGTGTCTGTCTGCTGAATTATAGACAGGATGGCACAACCTTCTGGAATCTCTTCTTCGTGGCTGGACTCAGAGAT<br>TCTAAGGGCAATATTGTCAACTACGTCGGAGTGCAGTCAAAGGTGAGCGAAGATTATGCCAAGCTGCTGGTCAACGAGCA<br>GAACATTGAGTACAAAGGTGTGCGCACCAGTAACATGCTGCGCAGAAAGCCCGGGTAA |
| >pOVC1_casp9_CrPH-LOV | ACCATGGCCACCGGGGGATTTGGTGATGTCGGTGCTCTTGAGAGTTTGAGGGGAAATGCAGATTTGGCTTACATCCTGAG<br>CATGGAGCCCTGTGGCCACTGCCTCATTATCAACAATGTGAACTTCTGCCGTGAGTCCGGGCTCCGCACCCGCACTGGCT<br>CCAACATCGACTGTGAGAAGTTGCGGCGTCGCTTCTCCTCGCTGCATTTCATGGTGGAGGTGAAGGGCGACCTGACTGCC<br>AAGAAAATGGTGCTGGCTTTGCTGGAGCTGGCGCGGCAGGACCACGGTGCTCTGGACTGCTGCGTGGTGGTCATTCTCTC<br>TCACGGCTGTCAGGCCAGCCACCTGCAGTTCCCAGGGGCTGTCTACGGCACAGATGGATGCCCTGTGTCGGTCGAGAAGA<br>TTGTGAACATCTTCAATGGGACCAGCTGCCCCAGCCTGGGAGGGAAGCCCAAGCTCTTTTTCATCCAGGCCTGTGGTGGG<br>GAGCAGAAAGACCATGGGTTTGAGGTGGCCTCCACTTCCCCTGAAGACGAGTCCCCTGGCAGTAACCCCGAGCCAGATGC<br>CACCCCGTTCCAGGAAGGTTTGAGGACCTTCGACCAGCTGGACGCCATATCTAGTTTGCCCACACCCAGTGACATCTTTG<br>TGTCCTACTCTACTTTCCCAGGTTTTGTTTCCTGGAGGGACCCCAAGAGTGGCTCCTGGTACGTTGAGACCCTGGACGAC<br>ATCTTTGAGCAGTGGGCTCACTCTGAAGACCTGCAGTCCCTCCTGCTTAGGGTCGCTAATGCTGTTTCGGTGAAAGGGAT<br>TTATAAACAGATGCCTGGTTGCTTTAATTTCCTCCGGAAAAAACTTTTCTTTAAAACATCACCCGGTGAATTCTCAGGCT<br>CTGGATCCCCCGGTGCAGGACTCAGACATACATTTGTGGTGGCTGATGCAACACTCCCTGATTGCCCACTGGTCTATGCA<br>AGTGAGGGCTTCTACGCAATGACCGGATATGGACCTGACGAAGTGCTGGGTCACAACTGTAGGTTTCTGCAGGGTGAGGG<br>AACTGACCCCAAGGAAGTGCAGAAAATTCGCGACGCCATCAAGAAGGGTGAGGCTTGTAGTGTGCGCCTCCTGAACTATC<br>GGAAGGACGGCACTCCCTTCTGGAACCTGCTGACAGTCACCCCAATTAAAACCCCTGATGGCCGCGTGTCCAAGTTTGTC<br>GGCGTGCAGGTGGATGTTACCTCCAAGACTGAAGGGAAAGCCCTGGCCCCCGGGTAA |
| >pOVC1_casp9_NcVVD-LOV | ACCATGGCCACCGGGGGATTTGGTGATGTCGGTGCTCTTGAGAGTTTGAGGGGAAATGCAGATTTGGCTTACATCCTGAG<br>CATGGAGCCCTGTGGCCACTGCCTCATTATCAACAATGTGAACTTCTGCCGTGAGTCCGGGCTCCGCACCCGCACTGGCT<br>CCAACATCGACTGTGAGAAGTTGCGGCGTCGCTTCTCCTCGCTGCATTTCATGGTGGAGGTGAAGGGCGACCTGACTGCC |

| | |
|---|---|
| | AAGAAAATGGTGCTGGCTTTGCTGGAGCTGGCGCGGCAGGACCACGGTGCTCTGGACTGCTGCGTGGTGGTCATTCTCTC<br>TCACGGCTGTCAGGCCAGCCACCTGCAGTTCCCAGGGGCTGTCTACGGCACAGATGGATGCCCTGTGTCGGTCGAGAAGA<br>TTGTGAACATCTTCAATGGGACCAGCTGCCCCAGCCTGGGAGGGAAGCCCAAGCTCTTTTTCATCCAGGCCTGTGGTGGG<br>GAGCAGAAAGACCATGGGTTTGAGGTGGCCTCCACTTCCCCTGAAGACGAGTCCCCTGGCAGTAACCCCGAGCCAGATGC<br>CACCCCGTTCCAGGAAGGTTTGAGGACCTTCGACCAGCTGGACGCCATATCTAGTTTGCCCACACCCAGTGACATCTTTG<br>TGTCCTACTCTACTTTCCCAGGTTTTGTTTCCTGGAGGGACCCCAAGAGTGGCTCCTGGTACGTTGAGACCCTGGACGAC<br>ATCTTTGAGCAGTGGGCTCACTCTGAAGACCTGCAGTCCCTCCTGCTTAGGGTCGCTAATGCTGTTTCGGTGAAAGGGAT<br>TTATAAACAGATGCCTGGTTGCTTTAATTTCCTCCGGAAAAAACTTTTCTTTAAAACATCACCCGGTGAATTCTCAGGCT<br>CTGGATCCCCCGGGCACACTCTCTACGCCCCAGGCGGGTACGATATTATGGGCTGGCTGATCCAGATCATGAACAGGCCC<br>AATCCCCAGGTCGAGCTGGGACCCGTGGATACTTCATGTGCACTGATACTGTGCGACCTGAAGCAGAAGGATACACCTAT<br>AGTTTACGCTTCAGAAGCCTTTCTGTACATGACAGGGTATTCTAACGCCGAGGTGCTGGGGAGGAACTGTAGGTTCCTCC<br>AGAGTCCCGATGGTATGGTGAAACCTAAGAGTACTCGCAAATATGTGGATAGCAATACTATTAACACCATGAGGAAAGCC<br>ATCGACAGAAACGCAGAAGTTCAGGTGGAAGTGGTGAACTTTAAGAAGAACGGCCAGCGGTTCGTGAACTTTCTCACAAT<br>GATTCCAGTGCGGGACGAAACCGGGGAGTACCGGTACAGCATGGGTTTTCAGTGCGAAACCGAACCCGGGTAA |
| >pOVC1_casp9_Rs LP-LOV | ACCATGGCCACCGGGGGATTTGGTGATGTCGGTGCTCTTGAGAGTTTGAGGGGAAATGCAGATTTGGCTTACATCCTGAG<br>CATGGAGCCCTGTGGCCACTGCCTCATTATCAACAATGTGAACTTCTGCCGTGAGTCCGGGCTCCGCACCCGCACTGGCT<br>CCAACATCGACTGTGAGAAGTTGCGGCGTCGCTTCTCCTCGCTGCATTTCATGGTGGAGGTGAAGGGCGACCTGACTGCC<br>AAGAAAATGGTGCTGGCTTTGCTGGAGCTGGCGCGGCAGGACCACGGTGCTCTGGACTGCTGCGTGGTGGTCATTCTCTC<br>TCACGGCTGTCAGGCCAGCCACCTGCAGTTCCCAGGGGCTGTCTACGGCACAGATGGATGCCCTGTGTCGGTCGAGAAGA<br>TTGTGAACATCTTCAATGGGACCAGCTGCCCCAGCCTGGGAGGGAAGCCCAAGCTCTTTTTCATCCAGGCCTGTGGTGGG<br>GAGCAGAAAGACCATGGGTTTGAGGTGGCCTCCACTTCCCCTGAAGACGAGTCCCCTGGCAGTAACCCCGAGCCAGATGC<br>CACCCCGTTCCAGGAAGGTTTGAGGACCTTCGACCAGCTGGACGCCATATCTAGTTTGCCCACACCCAGTGACATCTTTG<br>TGTCCTACTCTACTTTCCCAGGTTTTGTTTCCTGGAGGGACCCCAAGAGTGGCTCCTGGTACGTTGAGACCCTGGACGAC<br>ATCTTTGAGCAGTGGGCTCACTCTGAAGACCTGCAGTCCCTCCTGCTTAGGGTCGCTAATGCTGTTTCGGTGAAAGGGAT<br>TTATAAACAGATGCCTGGTTGCTTTAATTTCCTCCGGAAAAAACTTTTCTTTAAAACATCACCCGGTGAATTCTCAGGCT<br>CTGGATCCCCCGGTGCCATGGATCAGAAGCAGTTTGAGAAGATTAGAGCTGTGTTTGACAGGTCAGGGGTCGCACTGACC<br>CTCGTTGACATGTCCCTGCCAGAGCAACCCCTGGTGCTCGCCAACCCTCCATTTCTGAGAATGACTGGCTATACTGAGGG<br>CCAGATCCTGGGATTCAACTGCAGATTTCTCCAGAGAGGCGACGAAAATGCTCAGGCACGGGCTGACATCAGAGATGCCC<br>TCAAGCTCGGAAGGGAGCTCCAGGTGGTCCTCCGCAATTACAGAGCCAACGATGAACCATTTGACAATCTGCTGTTCCTG<br>CACCCTGTCGGTGGCAGACCCGACGCTCCTGACTACTTCCTCGGTTCTCAGTTCGAGCTGGGTAGAAGCGGAAATAGCGA<br>AGAGGCAGCCGCAGCTGGACACGCAGGGGCACTGACTGGGGAGCTCGCCAGAATAGGAACTGTGGCTGCTCGGCTCGAAA<br>TGGACAGTCGGAGACATCTGGCACAAGCTGCTGCAGCCCTGGTGAGGGCCTGGGAAAGAAGGGGTCCCGGGTAA |
| >pOVC1_casp9_At CRY2-PHR | ACCATGGCCACCGGGGGATTTGGTGATGTCGGTGCTCTTGAGAGTTTGAGGGGAAATGCAGATTTGGCTTACATCCTGAG<br>CATGGAGCCCTGTGGCCACTGCCTCATTATCAACAATGTGAACTTCTGCCGTGAGTCCGGGCTCCGCACCCGCACTGGCT<br>CCAACATCGACTGTGAGAAGTTGCGGCGTCGCTTCTCCTCGCTGCATTTCATGGTGGAGGTGAAGGGCGACCTGACTGCC<br>AAGAAAATGGTGCTGGCTTTGCTGGAGCTGGCGCGGCAGGACCACGGTGCTCTGGACTGCTGCGTGGTGGTCATTCTCTC<br>TCACGGCTGTCAGGCCAGCCACCTGCAGTTCCCAGGGGCTGTCTACGGCACAGATGGATGCCCTGTGTCGGTCGAGAAGA<br>TTGTGAACATCTTCAATGGGACCAGCTGCCCCAGCCTGGGAGGGAAGCCCAAGCTCTTTTTCATCCAGGCCTGTGGTGGG<br>GAGCAGAAAGACCATGGGTTTGAGGTGGCCTCCACTTCCCCTGAAGACGAGTCCCCTGGCAGTAACCCCGAGCCAGATGC<br>CACCCCGTTCCAGGAAGGTTTGAGGACCTTCGACCAGCTGGACGCCATATCTAGTTTGCCCACACCCAGTGACATCTTTG<br>TGTCCTACTCTACTTTCCCAGGTTTTGTTTCCTGGAGGGACCCCAAGAGTGGCTCCTGGTACGTTGAGACCCTGGACGAC<br>ATCTTTGAGCAGTGGGCTCACTCTGAAGACCTGCAGTCCCTCCTGCTTAGGGTCGCTAATGCTGTTTCGGTGAAAGGGAT<br>TTATAAACAGATGCCTGGTTGCTTTAATTTCCTCCGGAAAAAACTTTTCTTTAAAACATCACCCGGTGAATTCTCAGGCT<br>CTGGATCCCCCGGTAAGATGGACAAGAAAACCATAGTCTGGTTTCGCCGCGATTTGAGGATAGAGGATAACCCTGCGTTG<br>GCCGCAGCGGCCCACGAAGGCAGCGTGTTCCCCGTGTTCATATGGTGCCCAGAGGAGGAAGGCCAGTTCTACCCAGGTCG<br>CGCTAGTCGCTGGTGGATGAAACAGTCTCTCGCACATCTCTCCCAATCTTTGAAAGCTCTCGGGTCTGACCTTACGCTGA<br>TTAAGACCCACAATACTATTAGTGCAATCTTGGACTGCATCCGCGTTACCGGCGCGACCAAAGTGGTGTTTAATCATCTG<br>TATGACCCGGTAAGTCTGGTACGCGACCATACAGTTAAGGAGAAGTTGGTTGAAAGGGGAATTTCAGTACAGAGTTATAA<br>TGGAGACCTCCTCTACGAACCTTGGGAAATATACTGTGAGAAGGGAAAGCCATTTACATCATTTAACTCATACTGGAAAA<br>AGTGTCTTGACATGAGCATAGAGTCTGTCATGTTGCCCCCGCCGTGGCGGCTTATGCCCATCACCGCGGCCGCCGAAGCT<br>ATCTGGGCGTGTAGTATTGAAGAACTTGGCTTGGAGAACGAAGCAGAAAAACCTAGCAATGCACTGTTGACGCGCGCCTG<br>GTCCCCTGGGTGGAGTAACGCAGATAAATTGCTTAACGAGTTCATCGAAAAGCAACTTATCGACTACGCTAAGAATAGCA<br>AAAAAGTGGTGGGCAATTCAACCTCCCTCCTGTCTCCGTACTTGCATTTCGGCGAGATTAGCGTGCGGCACGTCTTCCAA<br>TGCGCCCGAATGAAACAAATAATCTGGGCGCGGGATAAAAACAGTGAGGGTGAAGAAAGCGCAGACTTGTTCCTCCGAGG<br>AATCGGTCTGCGAGAATATAGTCGCTACATTTGTTTCAATTTTCCCTTTACGCATGAGCAGAGCCTCCTTAGTCACTTGC<br>GATTCTTTCCTTGGGACGCAGATGTTGACAAATTTAAAGCATGGCGGCAAGGTAGAACAGGCTACCCATTGGTAGATGCG<br>GGTATGCGAGAACTCTGGGCCACGGGGTGGATGCATAACCGAATCAGGGTAATAGTAAGTAGTTTCGCAGTTAAGTTTCT<br>TTTGCTTCCATGGAAGTGGGGGATGAAGTATTTCTGGGACACTTTGCTCGATGCGGATCTTGAATGCGATATATTGGGTT<br>GGCAATATATTTCCGGGTCAATCCCTGACGGCCATGAGCTGGACAGATTGGACAATCCTGCGCTCCAGGGTGCGAAATAC<br>GATCCCGAAGGTGAATACATTAGACAATGGCTTCCAGAACTTGCCAGGTTGCCCACGGAATGGATTCACCACCCATGGGA<br>CGCCCCTCTTACGGTTTTGAAGGCGAGTGGTGTGGAGCTCGGTACCAATTACGCAAAACCGATTGTTGACATTGATACCG<br>CGCGGGAGTTGCTGGCTAAAGCCATTTCACGAACCCGGAAAGCCCAAATCATGATTGGTGCTGCAACCGGGTAA |
| >pOVC1_casp9_Hs FKBP | ACCATGGCCACCGGGGGATTTGGTGATGTCGGTGCTCTTGAGAGTTTGAGGGGAAATGCAGATTTGGCTTACATCCTGAG<br>CATGGAGCCCTGTGGCCACTGCCTCATTATCAACAATGTGAACTTCTGCCGTGAGTCCGGGCTCCGCACCCGCACTGGCT<br>CCAACATCGACTGTGAGAAGTTGCGGCGTCGCTTCTCCTCGCTGCATTTCATGGTGGAGGTGAAGGGCGACCTGACTGCC<br>AAGAAAATGGTGCTGGCTTTGCTGGAGCTGGCGCGGCAGGACCACGGTGCTCTGGACTGCTGCGTGGTGGTCATTCTCTC<br>TCACGGCTGTCAGGCCAGCCACCTGCAGTTCCCAGGGGCTGTCTACGGCACAGATGGATGCCCTGTGTCGGTCGAGAAGA<br>TTGTGAACATCTTCAATGGGACCAGCTGCCCCAGCCTGGGAGGGAAGCCCAAGCTCTTTTTCATCCAGGCCTGTGGTGGG<br>GAGCAGAAAGACCATGGGTTTGAGGTGGCCTCCACTTCCCCTGAAGACGAGTCCCCTGGCAGTAACCCCGAGCCAGATGC<br>CACCCCGTTCCAGGAAGGTTTGAGGACCTTCGACCAGCTGGACGCCATATCTAGTTTGCCCACACCCAGTGACATCTTTG |

| | |
|---|---|
| | TGTCCTACTCTACTTTCCCAGGTTTTGTTTCCTGGAGGGACCCCAAGAGTGGCTCCTGGTACGTTGAGACCCTGGACGAC<br>ATCTTTGAGCAGTGGGCTCACTCTGAAGACCTGCAGTCCCTCCTGCTTAGGGTCGCTAATGCTGTTTCGGTGAAAGGGAT<br>TTATAAACAGATGCCTGGTTGCTTTAATTTCCTCCGGAAAAAACTTTTCTTTAAAACATCACCCGGTGAATTCTCAGGCT<br>CTGGATCCCCCGGTAAACTGGAAGTCGAGGGAGTGCAGGTGGAGACTATCTCCCCAGGAGACGGGCGCACCTTCCCCAAG<br>CGCGGCCAGACCTGCGTGGTGCACTACACCGGGATGCTTGAAGATGGAAAGAAAGTTGATTCCTCCTGGGACAGAAACAA<br>GCCCTTTAAGTTTATGCTAGGCAAGCAGGAGGTGATCCGAGGCTGGGAAGAAGGGGTTGCCCAGATGAGTGTGGGTCAGA<br>GAGCCAAACTGACTATATCTCCAGATTATGCCTATGGTGCCACTGGGCACCCAGGCATCATCCCACCACATGCCACTCTC<br>GTCTTCGATGTGGAGCTTCTAAAACTGGAATCTGGCGGTACCGGGTAA |
| >pOVC1_VfAU1-LOV-HA-P2a-myc_casp9 | ACCATGGCCACCGGTCCTGACTACAGTCTCGTGAAGGCTCTGCAAATGGCACAACAGAATTTTGTCATTACAGACGCCTC<br>CCTCCCAGACAACCCTATCGTCTACGCCAGTAGAGGGTTTCTGACACTGACAGGCTATTCTCTCGACCAGATCCTGGGCA<br>GGAACTGCAGGTTTCTGCAAGGGCCAGAAACAGACCCAAGAGCTGTGGATAAGATCAGGAATGCCATCACCAAAGGCGTT<br>GATACCAGTGTCTGTCTGCTGAATTATAGACAGGATGGCACAACCTTCTGGAATCTCTTCTTCGTGGCTGGACTCAGAGA<br>TTCTAAGGGCAATATTGTCAACTACGTCGGAGTGCAGTCAAAGGTGAGCGAAGATTATGCCAAGCTGCTGGTCAACGAGC<br>AGAACATTGAGTACAAAGGTGTGCGCACCAGTAACATGCTGCGCAGAAAGCCCGGTGAATTCTACCCATACGATGTTCCA<br>GATTACGCTGCGACCAACTTTAGCCTGCTGAAACAGGCGGGCGATGTGGAAGAAAACCCGGGCCCGGAACAAAAACTCAT<br>CTCAGAAGAGGATCTGGGATCCCCCGGGGGATTTGGTGATGTCGGTGCTCTTGAGAGTTTGAGGGGAAATGCAGATTTGG<br>CTTACATCCTGAGCATGGAGCCCTGTGGCCACTGCCTCATTATCAACAATGTGAACTTCTGCCGTGAGTCCGGGCTCCGC<br>ACCCGCACTGGCTCCAACATCGACTGTGAGAAGTTGCGGCGTCGCTTCTCCTCGCTGCATTTCATGGTGGAGGTGAAGGG<br>CGACCTGACTGCCAAGAAAATGGTGCTGGCTTTGCTGGAGCTGGCGCGGCAGGACCACGGTGCTCTGGACTGCTGCGTGG<br>TGGTCATTCTCTCTCACGGCTGTCAGGCCAGCCACCTGCAGTTCCCAGGGGCTGTCTACGGCACAGATGGATGCCCTGTG<br>TCGGTCGAGAAGATTGTGAACATCTTCAATGGGACCAGCTGCCCCAGCCTGGGAGGGAAGCCCAAGCTCTTTTTCATCCA<br>GGCCTGTGGTGGGGAGCAGAAAGACCATGGGTTTGAGGTGGCCTCCACTTCCCCTGAAGACGAGTCCCCTGGCAGTAACC<br>CCGAGCCAGATGCCACCCCGTTCCAGGAAGGTTTGAGGACCTTCGACCAGCTGGACGCCATATCTAGTTTGCCCACACCC<br>AGTGACATCTTTGTGTCCTACTCTACTTTCCCAGGTTTTGTTTCCTGGAGGGACCCCAAGAGTGGCTCCTGGTACGTTGA<br>GACCCTGGACGACATCTTTGAGCAGTGGGCTCACTCTGAAGACCTGCAGTCCCTCCTGCTTAGGGTCGCTAATGCTGTTT<br>CGGTGAAAGGGATTTATAAACAGATGCCTGGTTGCTTTAATTTCCTCCGGAAAAAACTTTTCTTTAAAACATCACCCGGG<br>TAA |

### References (Supplementary Material)


1. Grusch, M., Schelch, K., Riedler, R., Reichhart, E., Differ, C., Berger, W., Ingles-Prieto, A. and Janovjak, H. (2014) Spatio-temporally precise activation of engineered receptor tyrosine kinases by light. *EMBO J*, **33**, 1713-1726.
2. Wang, X., Chen, X. and Yang, Y. (2012) Spatiotemporal control of gene expression by a light-switchable transgene system. *Nat Methods*, **9**, 266-269.
3. Conrad, K.S., Bilwes, A.M. and Crane, B.R. (2013) Light-induced subunit dissociation by a light-oxygen-voltage domain photoreceptor from Rhodobacter sphaeroides. *Biochemistry*, **52**, 378-391.
4. Guntas, G., Hallett, R.A., Zimmerman, S.P., Williams, T., Yumerefendi, H., Bear, J.E. and Kuhlman, B. (2015) Engineering an improved light-induced dimer (iLID) for controlling the localization and activity of signaling proteins. *Proc Natl Acad Sci U S A*, **112**, 112-117.
5. Strickland, D., Lin, Y., Wagner, E., Hope, C.M., Zayner, J., Antoniou, C., Sosnick, T.R., Weiss, E.L. and Glotzer, M. (2012) TULIPs: tunable, light-controlled interacting protein tags for cell biology. *Nat Methods*, **9**, 379-384.
6. Bugaj, L.J., Choksi, A.T., Mesuda, C.K., Kane, R.S. and Schaffer, D.V. (2013) Optogenetic protein clustering and signaling activation in mammalian cells. *Nat Methods*, **10**, 249-252.
7. Kennedy, M.J., Hughes, R.M., Peteya, L.A., Schwartz, J.W., Ehlers, M.D. and Tucker, C.L. (2010) Rapid blue-light-mediated induction of protein interactions in living cells. *Nat Methods*, **7**, 973-975.
8. Kainrath, S., Stadler, M., Reichhart, E., Distel, M. and Janovjak, H. (2017) Green-Light-Induced Inactivation of Receptor Signaling Using Cobalamin-Binding Domains. *Angew Chem Int Ed Engl*, **56**, 4608-4611.
9. Reichhart, E., Ingles-Prieto, A., Tichy, A.M., McKenzie, C. and Janovjak, H. (2016) A phytochrome sensory domain permits receptor activation by red light. *Angew Chem Int Ed Engl*, **55**, 6339-6342.
10. Levskaya, A., Weiner, O.D., Lim, W.A. and Voigt, C.A. (2009) Spatiotemporal control of cell signalling using a light-switchable protein interaction. *Nature*, **461**, 997-1001.
